# BCL11B predetermines a persister state in breast cancer that is reversed by TNF*α*

**DOI:** 10.64898/2026.06.09.731253

**Authors:** Zhen Qi, Gunsagar S. Gulati, Angera H. Kuo, Shaheen S. Sikandar, William Hai Dang Ho, Dalong Qian, Frederick M. Dirbas, Aaron M. Newman, Shang Cai, Michael F. Clarke

## Abstract

The frequent development of chemoresistance in cancer presents a significant clinical challenge, yet the underlying causes for heterogeneous drug responses remain largely elusive. Here, we systematically assessed the cellular differentiation status of human breast cancer cells utilizing single-cell atlases and identified a distinct population of immature basal-like cancer cells marked by *BCL11B*. Notably, higher levels of *BCL11B*^+^ cancer cells are significantly associated with early relapse in breast cancer patients who received chemotherapy. Functioning as a central regulator, BCL11B delineates an immature cell state that preferentially transitions to a drug-resistant persister state during treatment through multiple pre-existing and adaptive drug-resistance programs. Importantly, the cytokine tumor necrosis factor alpha (TNFα) is revealed as a natural inhibitor of BCL11B and can directly reverse the emergence of chemoresistant persister cells. Therefore, we identify BCL11B as an unappreciated pre-determinant of drug response and a therapeutic target for a subset of breast cancer patients at high risk of developing chemoresistance.

## Introduction

Despite showing an initial response to chemotherapy, a multitude of tumors, including those in breast cancer, develop treatment resistance in an unpredictable manner, posing a significant challenge in cancer management (*1*). While genetic heterogeneity can contribute to drug resistance, it is increasingly recognized that non-genetic mechanisms play a key role in the process (*2*). It has been suggested that a subset of cancer cells can enter a reversible drug-tolerant persister state to acquire resistance during treatment (*3, 4*). However, the diverse responses of tumors to chemotherapy treatment observed in the clinic, which can range from complete to minimal responses (*5, 6*), suggest that the emergence of cancer persisters is complex and likely involves both primary and adaptive mechanisms that remain largely unexplored. Growing evidence suggests that tumors exhibit heterogeneity at both inter-tumor and intra-tumor levels and contain diverse cell lineages or epigenetic cell states with intrinsic variations in biological features (*7, 8*). This indicates that there might be tumor subpopulations with pre-existing fitness advantages that can preferentially adopt a chemoresistant persister state following treatment. Consistent with this concept, recent lineage-tracing studies in colorectal cancer and skin tumors provide direct evidence that certain pre-existing tumor subpopulations selectively survive drug treatment and drive tumor regeneration (*9–11*). Therefore, understanding the heterogeneity of cancer cell states and the presence of pre-existing drug-resistant populations is essential for identifying biomarkers that can predict drug response, particularly in breast cancer, where specific biomarkers are largely lacking. Moreover, devising effective therapeutic strategies to target cells with inherent resistance capacity remains challenging but could offer a promising opportunity to reverse the emergence of cancer persistence during treatment.

Breast cancer is a highly heterogeneous disease that encompasses multiple subtypes with different clinical outcomes (*12*). Breast cancer is categorized into three major subtypes based on the expression of two hormone receptors and the epidermal growth factor receptor HER2, and can also be stratified into several molecular subtypes according to gene expression profiling (*12*). Adding to the complexity, cancer cell populations in distinct cell states can co-exist in a single tumor, reflecting the normal mammary gland cellular hierarchy (*13*). The normal mammary epithelium comprises an inner luminal cell layer and an outer basal cell layer and harbors various cell states, including stem cells, luminal and basal progenitors, and mature luminal and myoepithelial cells (*14*). However, it remains unclear how the variability in initial cell states affects the drug responses in breast cancer.

Recent advances in single-cell RNA sequencing (scRNA-seq) technology have revolutionized our ability to study tumor cell heterogeneity and have uncovered previously unknown cell states and cellular trajectories (*15*). Here, we utilized single-cell atlases to evaluate the cellular differentiation states in human breast cancer and identified a previously unrecognized *BCL11B*^+^ immature cancer cell state that is inherently associated with chemotherapy resistance. As a zinc finger transcriptional factor, BCL11B plays a critical role in regulating fetal development and tissue homeostasis in multiple systems, including T cells, neural tissues, skin, and mammary gland (*16–20*). BCL11B has also been implicated in various malignancies in humans, exhibiting context-dependent functions (*21–24*). By combining human and mouse tumor models and single-cell profiling, we revealed here that in human breast cancer, BCL11B defines a pre-existing immature cancer subpopulation and confers the cells with a remarkable capacity to persist during chemotherapy by leveraging both inherent and adaptive mechanisms. Importantly, our findings indicate that BCL11B could potentially be used as a biomarker for predicting the risk of chemoresistance in breast cancer patients and as a druggable target to reverse the development of treatment resistance.

## Results

### BCL11B marks a distinct immature cancer cell population in human breast cancer

We have previously shown that the CytoTRACE model, a robust computational framework for predicting cellular differentiation states in scRNA-seq datasets, can be used to identify both normal stem cells and immature cells within cancers (*25*). To systematically investigate the immature cell states in breast cancer and reveal genes associated with tumor characteristics including therapy resistance, we applied CytoTRACE analysis to the whole cancer epithelial compartment in a single-cell atlas encompassing ER+, HER2+, and triple-negative breast cancer (TNBC) subtypes (*26*). Cancer cells from different patient samples were mainly grouped into separate clusters and exhibited varied levels of differentiation as inferred by CytoTRACE (**Fig. 1, A and B**, and fig. S1A), aligning with the concept of inter-tumor heterogeneity. The TNBC cells generally exhibited lower levels of differentiation compared to other cancer cells, while within each tumor sample, including TNBC, there were usually tumor subpopulations in a relatively less differentiated state (**Fig. 1B** and fig. S1A).

**Fig. 1.**
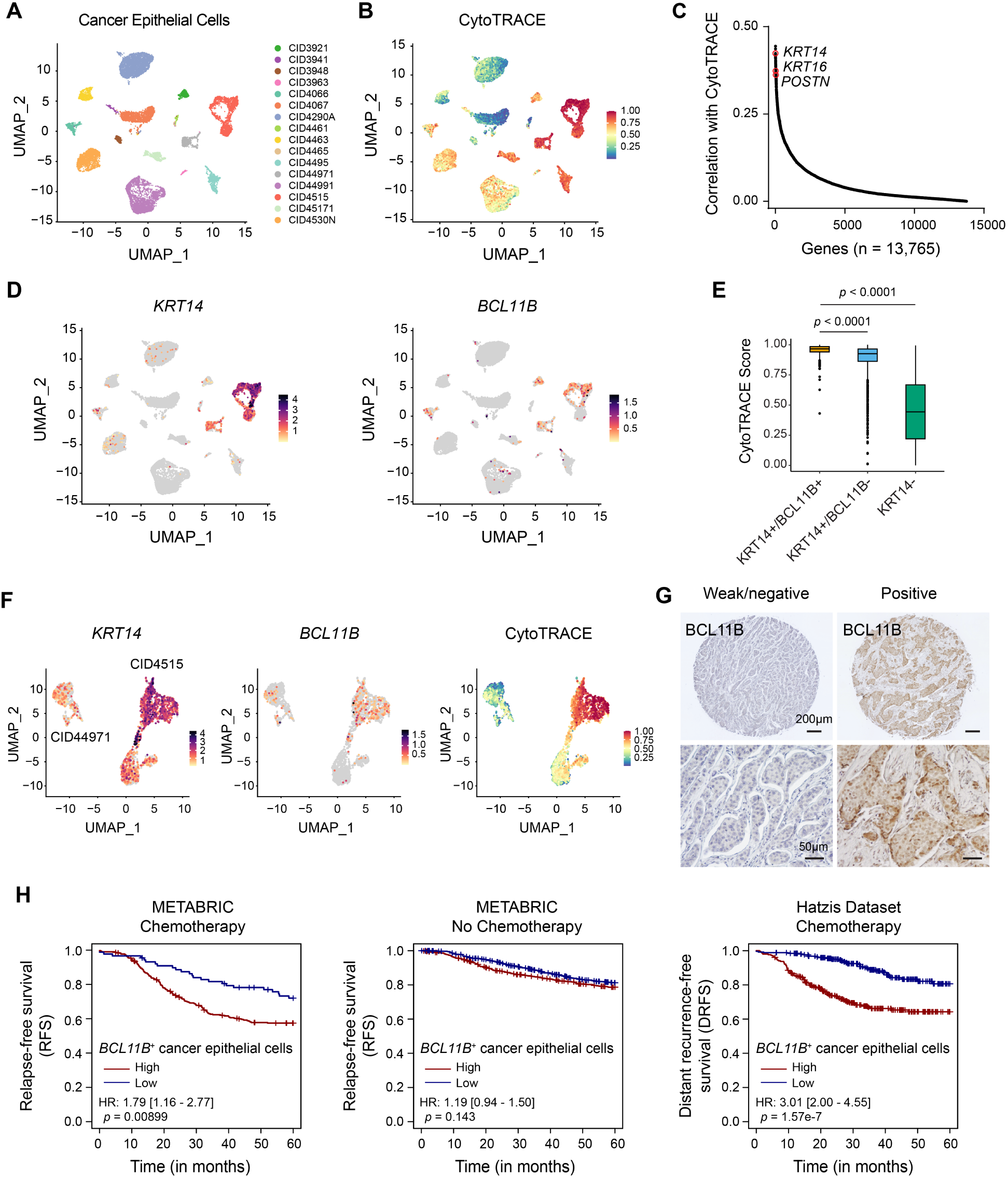
BCL11B marks an immature cancer cell state associated with chemotherapy resistance in human breast cancer. (**A**) UMAP representation of the cancer epithelial compartment from 16 human breast tumors (public scRNA-seq dataset available through the Broad Single-Cell portal and GEO: GSE176078; the metaplastic tumor samples were excluded and the IDC tumors were analyzed). (**B**) UMAP plot displaying the levels of CytoTRACE scores in the cancer epithelial compartment. Higher CytoTRACE scores indicate less differentiated cell states. (**C**) Plot showing the ranking of genes that are positively associated with CytoTRACE. Only genes with an expression percentage in all cancer cells below the mean percentage minus one standard deviation (<12.42%) were considered, in order to exclude genes with broader expression patterns and identify genes specific to less-differentiated cells. (**D**) UMAP plots displaying the expression patterns of *KRT14* and *BCL11B* in the cancer epithelial compartment. (**E**) Boxplot comparing the levels of CytoTRACE scores among the indicated groups in the entire cancer compartment. (**F**) UMAP plot displaying the expressions of *BCL11B* and *KRT14*, as well as the levels of CytoTRACE scores in *KRT14*-high cancer populations. Note that the *KRT14*-high cell clusters were derived from patients CID4515 and CID44971. (**G**) Representative images of BCL11B immunohistochemistry staining in human breast cancer samples. (**H**) Left and middle panel: Kaplan-Meier curve showing differences in relapse-free survival stratified by the median abundance of *BCL11B*^+^ cancer epithelial cells in the patients with or without chemotherapy treatment in the METABRIC cohort (n = 392, chemotherapy; n = 1,503, no chemotherapy); Right panel: Kaplan-Meier curve showing differences in distant recurrence-free survival stratified by the median abundance of *BCL11B*^+^ cancer epithelial cells in the Hatzis cohort (n = 508).

To explore common markers for the less differentiated cancer cells across the dataset, we ranked genes based on their correlation with CytoTRACE. Interestingly, *KRT14* was identified as one of the top three genes associated with less differentiated cells, along with other basal genes, such as *KRT16* and *POSTN*, ranking at the top of the list (**Fig. 1C** and table S1). While *KRT14* is typically seen as a marker for basal cells, it is also found in cells with basal-luminal intermediate features in the normal human breast (*27*). Here, *KRT14*^+^ cancer cells exhibited low levels of myo-contractile genes expressed by mature basal cells (fig. S1B), indicating an immature basal-like cell state distinct from the differentiated phenotype. Upon closer examination of the *KRT14*^+^ populations, we observed that there were subsets of cells with low CytoTRACE scores (fig. S1B). To further investigate the more immature subset of *KRT14*^+^ cells, we paired each of the top 1,000 genes identified by CytoTRACE with *KRT14* and examined the mean CytoTRACE scores associated with each gene and gene pair. This analysis yielded a list of candidate genes that not only predict less differentiated cells independently in the whole dataset, but also mark subsets of *KRT14*^+^ cells in the more immature cell state (fig. S1C). Remarkably, *BCL11B*, a marker gene for a minority quiescent stem cell population that we previously identified in the mouse mammary epithelium (*19*), emerged as one of the top 30 candidate genes (fig. S1C). Many of the other top candidate genes have been previously linked to increased tumorigenic ability and invasiveness in breast cancer, such as *PLA2G7*, *MSLN*, *MCAM*, and *CREB5* (fig. S1C) (*28–31*), validating the effectiveness of our approach.

We next focused on *BCL11B* and found that *BCL11B*^+^ cells represented distinct tumor subpopulations in a subset of breast tumors and were primarily located in the *KRT14*^+^ compartments (**Fig. 1D** and fig. S1D). These cells were present in all three major cancer subtypes, with a higher frequency observed in TNBC (fig. S1E). The *BCL11B*^+^ cells showed an obvious correlation with less differentiated cancer cells across the dataset, with the *KRT14*^+^/*BCL11B*^+^ cells exhibiting the highest CytoTRACE scores compared to other tumor subsets (**Fig. 1D and E**). To better visualize *BCL11B* marking the more immature subsets of *KRT14*^+^ cells, we focused on the distinct *KRT14*^+^ cancer populations revealed by the combined analysis and performed separate CytoTRACE analysis on these cells. Note that CytoTRACE outputs the relative ordering of input cells (relative scores) in each analysis. These *KRT14*^+^ cancer populations were derived from two tumor samples, both of which showed enriched *BCL11B* expression in their relatively more immature subsets (**Fig. 1F** and fig. S1, F and G). To validate our findings, we examined a different single-cell atlas of treatment-naive human invasive breast cancers including 12 TNBC, 14 ER+, and 3 HER2+ tumors (fig. S1, H and I). Consistently, *BCL11B* marked tumor subpopulations in a subset of patient samples spanning all three cancer subtypes and was more specifically enriched in cancer cells in less differentiated states (fig. S1, J and K).

To examine the protein expression pattern of BCL11B in cancer tissues, we conducted tissue microarray analysis of 188 breast tumor samples and observed a similar inter-tumor heterogeneity. We found prominent BCL11B expression in approximately 27% of ER-negative tumors and 19% of ER-positive tumors, with the highest expression levels mainly observed in the ER-negative subtype (**Fig. 1G** and fig. S2A). In addition, in the majority of BCL11B-positive cases, only a subset of tumor cells, typically located at the tumor-stromal border, displayed high expression of BCL11B (**Fig. 1G** and fig. S2A). Therefore, our analysis combining different single-cell atlases and tissue microarray analyses revealed that BCL11B^+^ cells represent a distinct tumor subpopulation in a group of breast cancer patients and exhibit a notable immature cell feature.

### *BCL11B*^+^ cancer cells are associated with chemotherapy resistance in breast cancer

Next, we sought to understand the clinical significance of *BCL11B* expression in breast cancer. To minimize the influence of non-cancerous expression of *BCL11B* in tumors, we utilized a deconvolution-based approach to reveal *BCL11B*-expressing cancer cells in bulk gene expression datasets. We defined transcriptional signatures for *BCL11B*^+^ cancer cells, *BCL11B*^-^ cancer cells, T cells, B cells and other niche cell types based on the single-cell atlas, and used CIBERSORTx to estimate the abundance of each cell type in individual samples in the METABRIC cohort (fig. S2B). Notably, we observed a strong association between higher levels of *BCL11B*^+^ cancer cells and early relapse (within 5 years) specifically in breast cancer patients who had received chemotherapy (**Fig. 1H**). No such correlation was found when chemotherapy was not given (**Fig. 1H**). The estimated levels of *BCL11B*^+^ tumor cells in breast cancer subtypes were comparable between the chemotherapy-treated and non-treated patients, indicating the robustness of the deconvolution method (fig. S2C). To validate this finding, we further analyzed the Hatzis breast cancer dataset (*32*), where over 93% of the patients received neoadjuvant chemotherapy. Consistently, higher abundances of *BCL11B*^+^ cancer cells were found to have a significant correlation with early distant recurrence in this chemotherapy-treated cohort (**Fig. 1H**). We also analyzed *KRT14* using a similar deconvolution method and observed that *KRT14* exhibited an inferior performance in predicting poor prognosis after chemotherapy when compared to BCL11B (fig. S2D), consistent with the more immature state of *BCL11B*^+^ cells within the *KRT14*^+^ population. Together, these data demonstrated a specific association between the elevated levels of *BCL11B*^+^ cancer cells and the high risk of developing chemotherapy resistance in human breast cancer.

### BCL11B defines a basal-like tumor subpopulation with pre-existing drug-resistance ability

HER2 has been identified as an important biomarker and treatment target for a subset of aggressive breast cancers, resulting in the development of highly successful targeted therapy (*33*). This prompted us to investigate the therapeutic significance of targeting BCL11B in addressing chemoresistance. We first experimentally examined whether the BCL11B-expressing cancer cells possess the ability to resist chemotherapeutic drugs. To this end, we utilized the widely used MMTV-PyMT (PyMT) transgenic mouse model, which has been shown to recapitulate many aspects of human breast cancer (*34*). Interestingly, despite being regarded as a luminal type of breast cancer, the primary PyMT tumor harbored a specific subset of KRT14^+^ basal-like cells, predominantly located at the tumor-stromal border and surrounding KRT8^+^ luminal tumor cells (**Fig. 2A**). This finding aligns with observations reported in recent studies (*35, 36*). Of note, BCL11B^+^ cells were found to be a notable subset of the KRT14^+^ cells in the tumor (**Fig. 2, B and C**). These results showed that BCL11B marks a distinctive tumor subpopulation in the PyMT tumor model, exhibiting basal-like features and resembling human BCL11B^+^ cancer cells.

**Fig. 2.**
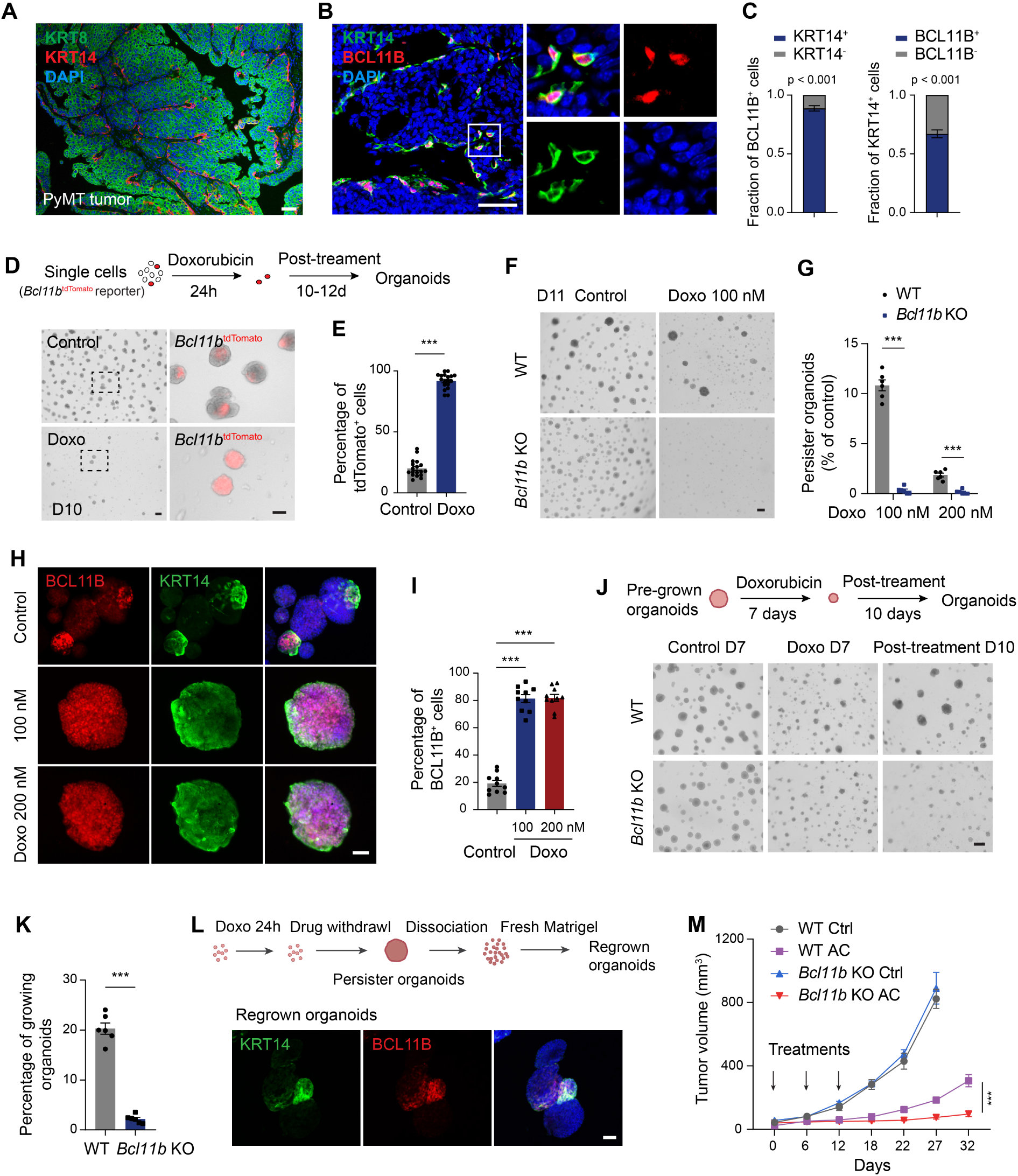
BCL11B^+^ cells represent a pre-existing drug-resistant cell population. (**A**) Immunofluorescence co-staining of KRT14/KRT8 in the primary PyMT tumor. (**B**) Representative immunostaining images of the primary tumor indicate the co-localization of BCL11B and KRT14. (**C**) Quantifications of the percentage of KRT14^+^/ KRT14^-^ cells within the BCL11B^+^ cells and the percentage of BCL11B^+^/ BCL11B^-^ cells within the KRT14^+^ population; n = 3 tumors. (**D**) Upper panel: schematic diagram showing the strategy of the *ex vivo* drug treatment assay. Lower panel: images showing the expression of tdTomato in the organoids from different groups on day 10. Scale bars, 200 μm (left) and 100 μm (right). (**E**) Quantification of the percentage of tdTomato^+^ cells in tumor organoids; n = 18 organoids from 3 experiments. (**F**) Single WT or *Bcl11b* KO tumor cells were cultured in the presence or absence of doxorubicin for 24 hours followed by drug washout. Representative images of tumor organoids on day 11 are shown. Scale bar, 200 μm. (**G**) Quantification of the frequency of drug-resistant persister organoids (% of control); n = 6 cell culture wells from 3 experiments. (**H**) Control organoids and drug-resistant organoids grown from the WT tumor cells shown in (**F**) were harvested and analyzed by immunostaining of KRT14 and BCL11B. Scale bar, 50 μm. (**I**) Percentage of BCL11B^+^ cells in tumor organoids under the indicated conditions; n = 10 organoids from 3 experiments. (**J**) Organoids were pre-grown for 3 days and then treated with doxorubicin (100 nM) for 7 days followed by drug washout. Representative images of tumor organoids are shown. Scale bar, 100 μm. (**K**) Percentage of actively growing organoids on day 17 in the indicated conditions; n = 6 cell culture wells from 3 experiments. (**L**) Drug-resistant organoids were dissociated into single cells and re-embedded in fresh Matrigel. Immunostaining of KRT14 and BCL11B in the regrown organoids was shown. Scale bar, 50 μm. (**M**) Tumor growth curves showing WT and *Bcl11b* KO tumor growth upon saline treatment, or combination treatment (AC, doxorubicin (1 mg/kg) + cyclophosphamide (60 mg/kg)). n = 6 mice per group, two-way ANOVA. Nuclei were counter-stained with DAPI (blue). Data are presented as mean ± SEM. *p < 0.05, **p < 0.01, ***p < 0.001.

To directly test whether the BCL11B^+^ cancer cell subpopulation possesses a pre-existing capability to resist treatment, we generated the *Bcl11b* reporter model by crossing the *Bcl11b*^tdTomato^ mice to MMTV-PyMT. *Bcl11b*^tdTomato^-positive cells represented a small subpopulation of tumor cells in the primary tumor as revealed by fluorescence-activated cell sorting (FACS) analysis (fig. S3A). We next established an *ex vivo* drug treatment assay starting with individual tumor cells. Briefly, single EpCAM^+^ tumor cells were sorted from PyMT-*Bcl11b*^tdTomato^ reporter tumors and cultured in Matrigel for 24 hours in the absence or presence of doxorubicin (Doxo), one of the most widely used chemotherapy agents for treating breast cancer (*37*). The drug was then washed out, and the cells were assayed for their capacity to form tumor organoids, a model system that has been shown to retain key properties of primary tumors (*38*) (**Fig. 2D**). In the vehicle-treated control group, PyMT tumor cells robustly grew into organoids within a short period of time (3-4 days) (**Fig. 2D**). However, following doxorubicin treatment, only a small fraction of cells survived and gradually developed into organoids which became evident after 7 days (**Fig. 2D**). Strikingly, when we analyzed the tdTomato fluorescence, we observed that the drug-resistant persister organoids were almost exclusively composed of *Bcl11b*^tdTomato^-positive cells, whereas only a minor subset of cells expressed tdTomato in the control organoids (**Fig. 2, D and E**). To explore whether *Bcl11b* is functionally required for the drug-resistant ability of this tumor population, we generated *Bcl11b* knockout (KO) tumor organoids by CRISPR/Cas9 (fig. S3B). *Bcl11b* deficiency did not cause an obvious change in the survival of the tumor organoids under normal conditions but led to a near-complete elimination of persister organoids following treatment (**Fig. 2, F and G**, and fig. S3C). Immunostaining analysis of the persister cells derived from WT tumor cells revealed that KRT14 and BCL11B double-positive cells were highly enriched in these organoids (**Fig. 2, H and I**). By contrast, the BCL11B^+^/KRT14^+^ tumor cells only constituted a subset of WT control organoids (**Fig. 2, H and I**). Together, these results reveal that BCL11B^+^ tumor cells constitute a pre-existing drug-tolerant population that can selectively persist during chemotherapy treatment.

To study the effects of prolonged treatment, we further administered doxorubicin to pre-grown WT or *Bcl11b*-deficient tumor organoids for 7 days (**Fig. 2J**). The majority of tumor organoids in both groups demonstrated a decrease in size following the treatment, although a small number of slow-growing organoids were observed in the WT group (**Fig. 2J**). Notably, upon termination of the treatment, a significant portion of the organoids in the WT group resumed growth, whereas the *Bcl11b*-deficient organoids were lost over time (**Fig. 2, J and K**). We also found that the absence of *Bcl11b* rendered tumor cells much more sensitive to paclitaxel (taxane), another drug commonly used to treat breast cancer (*37*), highlighting the multiple-drug resistance capacity of the BCL11B-expressing cells (fig. S3, D and E). We then asked whether the persister cells maintain a stable phenotype upon the discontinuation of drug treatment. To eliminate any potential negative impacts of residual drugs and to ensure any organoids were clonally derived from BCL11B^+^ cells, we dissociated the persister organoids into single cells and re-embedded them in fresh Matrigel. Interestingly, in the regrown organoids, the percentage of KRT14^+^/BCL11B^+^ cells dropped to a level similar to that of the pre-treatment organoids (**Fig. 2L** and fig. S3F), implying the differentiation of BCL11B^+^ cells. Moreover, the regrown organoids remained responsive to doxorubicin treatment due to the presence of drug-sensitive BCL11B^-^ cells (fig. S3, G and H). This indicates that BCL11B^+^ cells drive a reversible cancer persistence under drug treatment.

To test whether BCL11B expression enables tumor cells to establish drug-resistant ability, we reintroduced BCL11B expression in the *Bcl11b* knockout organoid cells (fig. S3I). We found that the restoration of BCL11B expression induced a *de novo* drug-resistance phenotype in these cells (fig. S3, J and K). To further confirm the role of BCL11B *in vivo*, we grew tumors from WT and *Bcl11b* knockout PyMT cells in mice and performed a drug treatment study. Consistent with our *in vitro* findings, knockout of *Bcl11b* dramatically increased the anti-tumor efficacy of either doxorubicin alone or the commonly used combination chemotherapy (AC, adriamycin (doxorubicin) combined with cyclophosphamide) (**Fig. 2M** and fig. S3L). This result also confirms that BCL11B confers resistance to multiple chemotherapeutic drugs that are frequently used in clinics. We also observed that drug treatment induced an obvious enrichment of BCL11B^+^ cells and KRT14^+^ cells in the WT residual tumor (fig. S3, M and N), consistent with the role of BCL11B in mediating treatment resistance. Overall, these *in vitro* and *in vivo* observations indicate that BCL11B shapes a pre-existing tumor population that is tolerant to multiple chemotherapeutic drugs and drives a reversible cancer persistence upon treatment.

### *The Bcl11b*^+^ cell state is intrinsically associated with drug-resistant properties

To investigate how *Bcl11b*^+^ cells are primed to adopt a drug-resistant state, we first performed bulk RNA-seq analysis on *Bcl11b*^+^ and *Bcl11b*^-^ cells in treatment-naïve tumors. As expected, basal genes such as *Col17a1*, *Krt14*, and *Krt5* were highly expressed by *Bcl11b*^+^ cells, while luminal markers were predominantly observed in *Bcl11b*^-^ cells (**Fig. 3A** and table S2). However, many of the top enriched genes in *Bcl11b*^+^ cells, such as *Dsc3*, *Dsg3*, and *Tgfbi*, have not been previously linked to mammary basal cells. Conversely, markers associated with a more mature myoepithelial phenotype, including *Actg2*, *Myh11*, *Myl9*, and *Mylk*, were found to be expressed at low levels in *Bcl11b*^+^ cells (**Fig. 3B**), consistent with our observations in human *BCL11B*^+^ cells. KEGG pathway analysis further uncovered an enrichment of genes involved in xenobiotic metabolism and glutathione (GSH) metabolism in the treatment-naïve *Bcl11b*^+^ cells, such as *Aldh3a1*, *Cyp26a1*, and *Mgst2* (**Fig. 3C and D**). Both pathways have been closely associated with detoxifying chemotherapeutic drugs in cancer (*39, 40*). Notably, one of the representative genes, the aldehyde dehydrogenase enzyme *Aldh3a1*, has been demonstrated in breast cancer to protect cells against various chemotherapeutic drugs including doxorubicin, paclitaxel, and cyclophosphamide (*41, 42*). By interrogating a recent single-cell transcriptome atlas spanning mouse mammary development stages (*43*), we also unexpectedly identified that *Bcl11b* and its associated gene program in cancer were specifically enriched in the rudimentary basal-like cells at the early postnatal stage (D5) (**Fig. 3E** and fig. S4, A-C). The majority of basal-like cells at this early postnatal stage lacked myoepithelial markers (**Fig. 3E**) and represented a precursor population in an intermediate developmental stage (*14, 44*). Interestingly, many of the xenobiotic detoxification genes also showed a specific enrichment in this early basal cell state (**Fig. 3E** and fig. S4C). Therefore, these results suggest that *Bcl11b*^+^ tumor cells mimic a precursor-like developmental state in cancer, which is inherently linked to a xenobiotic inactivation program that may play a physiological role in responding to external stimuli.

**Fig. 3.**
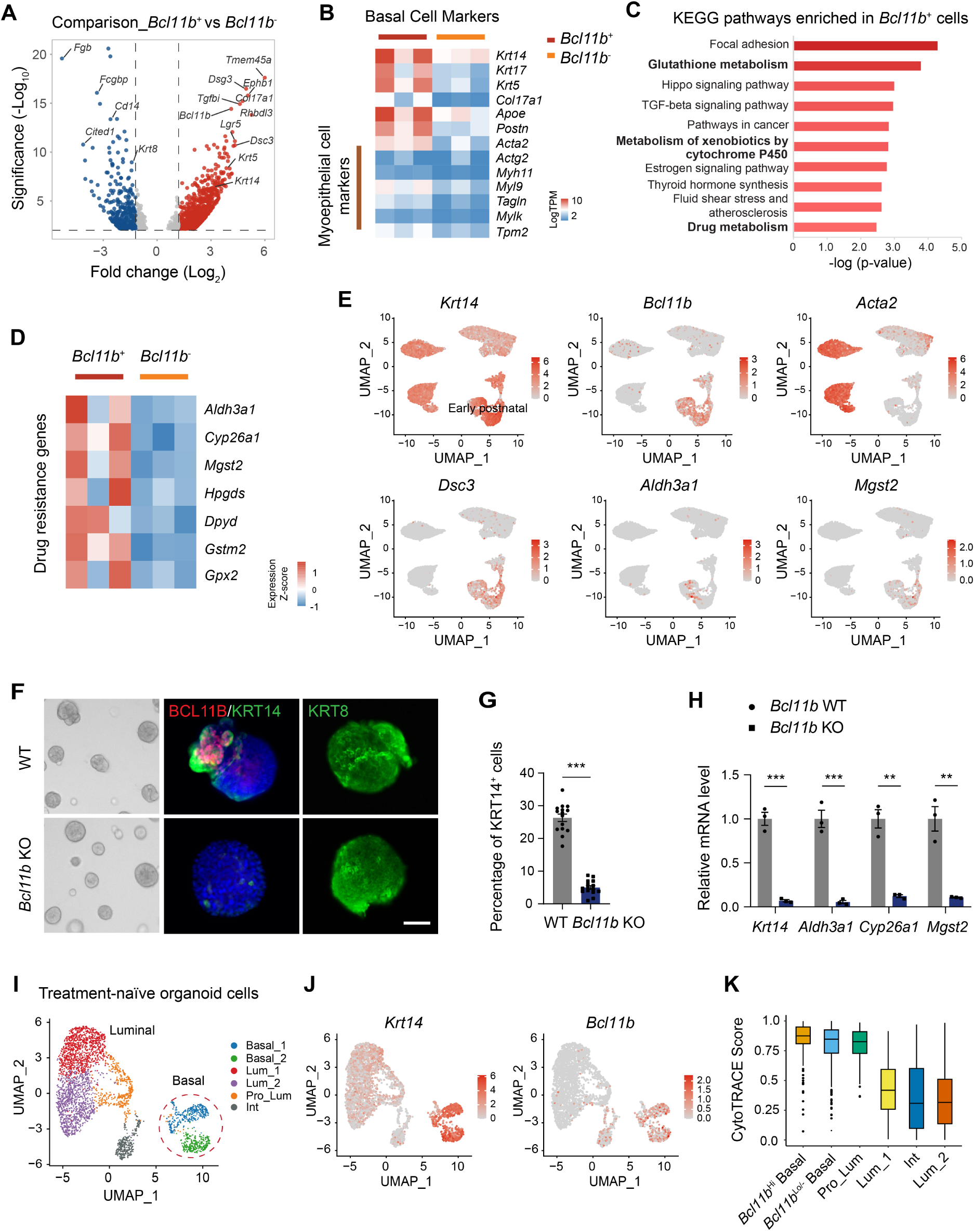
BCL11B functionally governs an immature basal-like cell state with inherent drug-resistance abilities. (**A**) Volcano plot showing genes differentially expressed between *Bcl11b*^+^ and *Bcl11b*^-^ PyMT tumor cells. (**B**) Heatmap visualization of basal marker genes and myoepithelial marker genes in the bulk RNA-seq dataset of *Bcl11b*^+^ and *Bcl11b*^-^ tumor cells. (**C**) KEGG pathway analysis of the upregulated genes in *Bcl11b*^+^ PyMT tumor cells. (**D**) Heatmap showing the expression levels of representative drug resistance genes in the bulk RNA-seq data of *Bcl11b*^+^ and *Bcl11b*^-^ cells. (**E**) UMAP plots displaying the expression patterns of indicated genes in the scRNA-seq dataset of mammary basal and basal-like cells spanning 4 developmental stages: embryonic, early postnatal, pre-puberty, and adult. (**F**) Left panel: bright-field images of wildtype (WT) and *Bcl11b* knockout (KO) tumor organoids; Middle panel: immunofluorescence co-staining of KRT14 and BCL11B in WT and *Bcl11b* KO tumor organoids; Right panel: Immunofluorescence staining of KRT8 in WT and *Bcl11b* KO tumor organoids. (**G**) Percentage of KRT14^+^ cells in WT and *Bcl11b* KO tumor organoids; n = 15 organoids from 3 experiments. (**H**) Real-time PCR analysis of indicated genes in WT and *Bcl11b* KO tumor cells with *Actb* as an internal control; n = 3 experiments. (**I**) UMAP representation of 3,377 vehicle-treated tumor organoid cells. The cells are colored by clusters. (**J**) UMAP plots displaying the expression patterns of indicated genes in the control organoid cells. (**K**) Boxplot showing the CytoTRACE scores of indicated tumor populations. Nuclei were counter-stained with DAPI (blue). Data are presented as mean ± SEM. *p < 0.05, **p < 0.01, ***p < 0.001.

Given that the deficiency of *Bcl11b* compromised the drug resistance ability, we questioned whether *Bcl11b* is functionally required for maintaining the immature basal-like cell state and its associated drug resistance program. Indeed, the loss of *Bcl11b* resulted in a remarkable decrease in the number of KRT14^+^ tumor cells, both *in vitro* and *in vivo*, while leaving the KRT8^+^ luminal cell lineage unaffected (**Fig. 3, F and G**, and fig. S4, D and E). The mRNA levels of *Krt14* and the drug resistance genes, including *Aldh3a1*, *Cyp26a1*, and *Mgst2*, were also significantly downregulated in tumor organoids after the knockout of *Bcl11b* (**Fig. 3H**). Transcriptomic analysis of WT and *Bcl11b* knockout tumor organoids further confirmed that both the basal gene program and the drug resistance program were impaired upon the deletion of *Bcl11b* (fig. S4F and Table S3). Together, these results suggest that *Bcl11b* controls an immature basal-like cell state that is inherently linked to a pre-existing drug detoxification gene program and exhibits selective advantages upon chemotherapy treatment.

### Single-cell transcriptomic characterization of the pre-treatment tumors

To better understand the early transcriptional changes in different tumor populations before and after treatment, we further conducted single-cell RNA sequencing (scRNA-seq) analysis on tumor organoids that were either untreated or treated with doxorubicin for 24 hours (**Fig. 3I** and fig. S5A). After quality filtering, we generated high-quality transcriptomic data for 6,764 tumor cells (3,377 treatment-naïve and 3,387 drug-treated cells), with a median of 3,700 genes detected per cell. We first analyzed the cell populations in the treatment-naïve tumor. UMAP analysis identified two major cell lineages (Basal and Luminal) divided into six cell clusters: Basal_1, Basal_2, Luminal_1 (Lum_1), Lum_2, Proliferating_Lum, and Intermediate cells (**Fig. 3I**). The basal-like tumor cells exhibited a distinct transcriptional profile from the luminal cells in the single-cell dataset (**Fig. 3J** and fig. S5B). The separation of basal-like cells into two closely related clusters was primarily driven by cell cycle heterogeneity, as a proportion of Basal_1 cells exhibited a moderate level of proliferation genes, while the Basal_2 cells were largely non-cycling (**Fig. 3I** and fig. S5, B and C). A highly proliferating luminal cell cluster was also evident as the cells strongly expressed the proliferation markers, such as *Mki67* (**Fig. 3I** and fig. S5, B and C). An intermediate cell population was identified as it co-expressed multiple luminal and basal cell markers (fig. S5, B and C).

The basal-like tumor cells were marked by *Krt14*, and a large proportion of the cells highly expressed *Bcl11b* (**Fig. 3J**). Immunostaining analysis of KRT14 and BCL11B protein expression in tumor organoids actually showed that the KRT14^+^ population comprises 62% BCL11B^High^ ^(Hi)^ cells, 28% BCL11B^Low^ ^(Lo)^ cells, and 10% BCL11B^-^ cells (fig. S5D). The BCL11B^Lo^ subset retains some BCL11B protein expression but it may have reduced levels of mRNA expression that could be underdetected in scRNA-seq. Therefore, we categorized the basal-like tumor cells in the single-cell dataset into *Bcl11b*^Hi^ cells (mRNA level > 0, 56%) and the remaining cells as *Bcl11b*^Lo/-^ cells. The latter group might still contain some *Bcl11b*-expressing cells, considering the immunostaining results and possible gene dropouts. Notably, the CytoTRACE analysis of the pre-treatment organoids revealed that the *Bcl11b*^Hi^ basal subset represented the least differentiated cells in the tumor organoids, and the basal-like tumor cells were generally less differentiated compared to luminal cells (**Fig. 3K**). In line with this, single *Bcl11b*^+^ cells can efficiently develop into tumor organoids comprising both KRT14^+^ basal-like cells and KRT8^+^ luminal cells (fig. S6A). This suggests that, similar to the normal murine mammary gland, the *Bcl11b*^+^ cells retain their developmental plasticity in the tumor. Moreover, *Bcl11b*-expressing cells demonstrated increased tumorigenicity compared to *Bcl11b*^-^ cells measured by *in vivo* transplantation and *ex vivo* tumor organoid forming assays (fig. S6, B-D). Microarray analysis of human breast tumor samples (*45*) confirmed that *BCL11B* expression was enriched in the tumorigenic-enriched population (CD44^+^CD24^-/low^) in human breast cancer (fig. S6E). Taken together, the transcriptomic analyses and functional assays reveal that BCL11B governs a unique immature basal-like cell state in the treatment-naïve tumor.

### *Bcl11b*^+^ tumor cells demonstrate adaptive resistance capacity during treatment

We next sought to understand how different tumor subpopulations respond to drugs. To this end, we performed a combined analysis of the pre- and post-treatment datasets. Of note, the untreated and drug-treated samples were processed and sequenced together, with no batch effect observed. We discovered that while the basal-like cells from different treatment conditions were largely overlapping on the UMAP plot, the luminal cells were separated into two distinct compartments that mainly corresponded to their treatment statuses (**Fig. 4, A and B**). Consistently, the untreated and drug-treated basal-like cells were almost evenly spread across the basal cell clusters, whereas drug-treated luminal cells were largely confined to two specific luminal clusters: Lum_3 and Lum_4 (**Fig. 4C**). Analysis of the cluster marker genes revealed that Lum_3 and Lum_4 cells displayed significant enrichment of p53, apoptosis, and interferon response-related pathways, while none of these pathways were enriched in basal-like tumor cells (**Fig. 4, D and E**, and fig. S7A). Moreover, the post-treatment luminal compartment showed a dramatic loss of *Ki67*^+^ proliferating cells (**Fig. 4, C and D**, and fig. S7B). By contrast, a considerable portion of basal cells retained *Ki67* expression after the treatment (fig. S**7**, B and C). These results suggest that basal-like cells were only moderately impacted by the drug treatment.

**Fig. 4.**
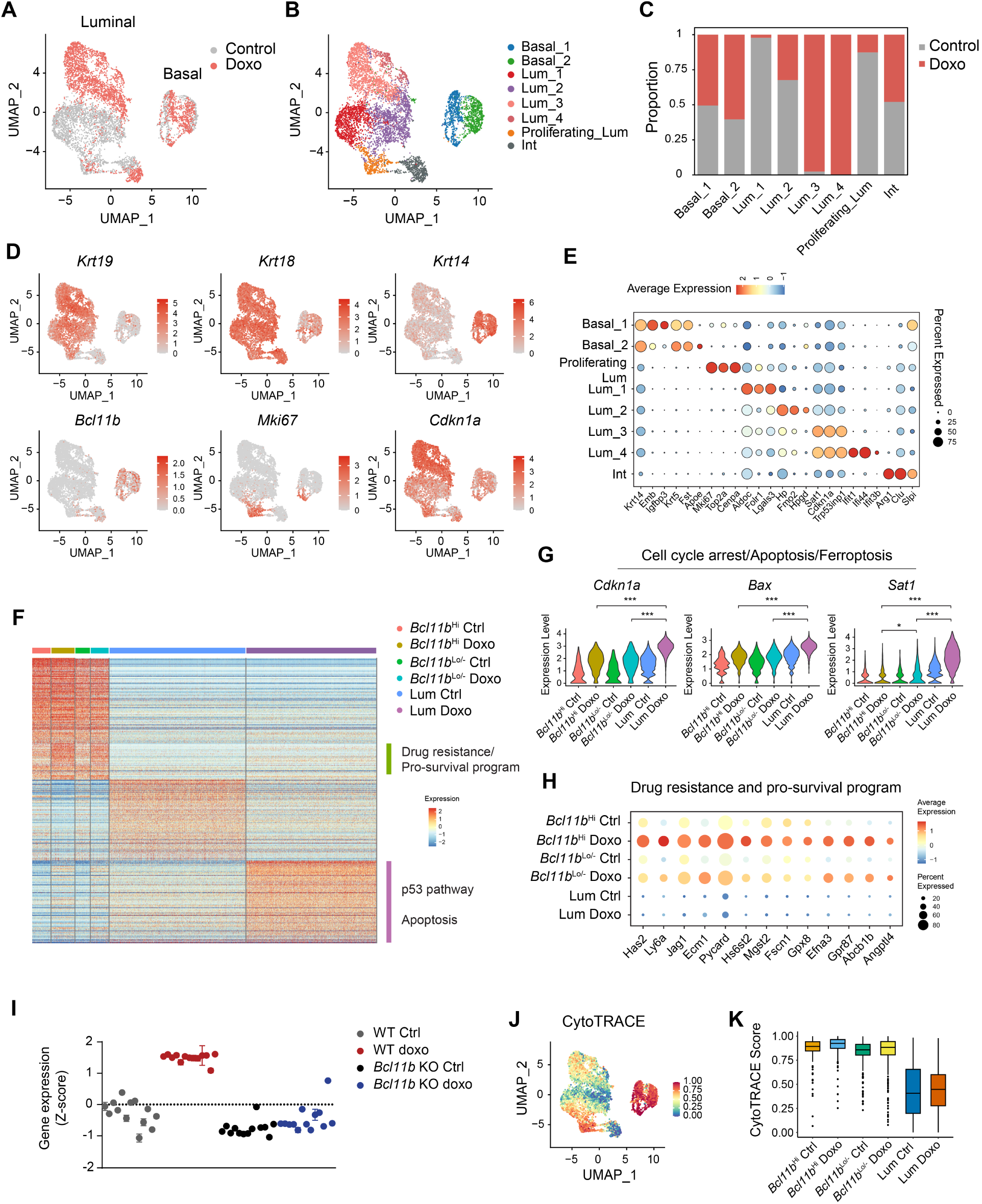
*Bcl11b*^+^ cancer cells demonstrate a notable capability to transcriptionally adapt to drug pressure. (**A**) UMAP representation of untreated and doxorubicin-treated tumor organoid cells. Cells are colored by treatment conditions. (**B**) Same UMAP plot as in (**A**) but colored by cell clusters. (**C**) The proportions of control and drug-treated cells across cell clusters. (**D**) UMAP plots showing the expression levels of selected marker genes. (**E**) Dotplot showing top marker genes for each cluster. (**F**) Heatmap showing the top 100 marker genes for the *Bcl11b*^Hi^ basal_Ctrl, *Bcl11b*^Hi^ basal_Doxo, *Bcl11b*^Lo/-^ basal_Ctrl, *Bcl11b*^Lo/-^ basal_Doxo, Lum_Ctrl, and Lum_Doxo. The top pathways enriched in the indicated clusters are shown. (**G**) Violin plots displaying the expression of representative Cell cycle arrest/Apoptosis genes and Drug Resistance/Survival/GSH metabolism genes across the six subsets. (**H**) Dotplot showing the expression levels of drug-resistance/pro-survival genes across the six indicated cell subsets. (**I**) Expression levels of the drug-resistance program in WT and *Bcl11b* KO tumor cells treated with vehicle or doxorubicin for 24 hours. Each dot represents an individual gene. Data are presented as mean ± SEM. (**J**) UMAP plot showing the levels of CytoTRACE scores in the combined dataset. (**K**) Boxplot comparing the levels of CytoTRACE scores among the indicated groups.

To gain a deeper insight into the transcriptional responses that occurred in different tumor subpopulations, we re-grouped the cells into six categories: *Bcl11b*^Hi^ basal_Ctrl, *Bcl11b*^Hi^ basal_Doxo, *Bcl11b*^Lo/-^ basal_Ctrl, *Bcl11b*^Lo/-^ basal_Doxo, Lum_Ctrl, and Lum_Doxo. As anticipated, pre- and post-treatment basal-like tumor cells were transcriptionally similar, while the drug treatment induced a drastic gene expression change in luminal cells featured by the activation of p53 and apoptosis pathways (**Fig. 4F**). Indeed, genes associated with cell cycle arrest, apoptosis and ferroptosis, such as *Cdkn1a*, *Btg2*, *Bax* and *Sat1* (*46–49*), were strongly induced in the luminal compartment following the treatment (**Fig. 4G**). We also noticed that a unique group of genes pre-existing in the basal compartment showed a specific upregulation in these cells upon treatment (**Fig. 4F**). The KEGG cancer pathways and GSH metabolism pathway are among the top-enriched pathways. As mentioned above, the GSH metabolism pathway contributes to the detoxification of drugs and reactive oxygen species (ROS) and has been found to be specifically enriched in chemoresistant squamous cell carcinoma stem cells and radiation-resistant breast cancer stem cells (*11, 50*). Moreover, many other cancer related genes within this gene group have previously been linked to various drug resistance and cell survival mechanisms, such as the activation of oncogenic signaling cascades (*Has2*, *Gpr87, Ecm1*, *Angptl4* (*51–55*)), heparan sulfate transferase-mediated drug resistance (*Hs6st2* (*56, 57*)), drug efflux (*Abcb1b* (*58*)), and enhanced anti-apoptosis functions (*Efna3* (*59*)) (**Fig. 4H**).

Despite technical issues causing the *Bcl11b*^Lo/-^ subset to potentially contain some *Bcl11b*-expressing cells and the fact that both subsets share a basal cell nature, the *Bcl11b*^Hi^ subset still displayed a stronger upregulation of the drug-resistance program compared to the *Bcl11b*^Lo/-^ tumor cells (**Fig. 4H**). Representative drug-resistance/pro-survival genes and GSH metabolism genes exhibited higher upregulation in the *Bcl11b*^Hi^ subset in response to treatment and low to no expression in luminal cells as shown in the Violin plots (fig. S7D). Importantly, the expression of the drug resistance gene program was predominantly dependent on *Bcl11b*, as drug treatment was unable to induce the majority of these genes in the *Bcl11b-*deficient tumor cells (**Fig. 4I** and table S4). Among the various drug-resistance pathways present or activated in *Bcl11b*^+^ cells, we specifically examined the drug detoxification program that involves both pre-existing and adaptive mechanisms, such as the drug metabolism gene *Aldh3a1* and the drug transporter *Abcb1b*. Consistent with our hypothesis, there was an obvious clearance of intracellular doxorubicin in BCL11B^+^ cells as opposed to BCL11B^-^ cells within 24 hours of administration (fig. S7, E and F). The uptake of doxorubicin in the cells was similar in both populations, supporting a drug elimination mechanism (fig. S7, E and F). Consistently, impairment of *Aldh3a1* or *Abcb1b* can significantly decrease the number of drug-resistant organoids following drug treatment (fig. S8, A-C). Moreover, the intracellular elimination of doxorubicin by BCL11B^+^ cells was effectively blocked when either *Aldh3a1* or *Abcb1b* was inhibited (fig. S8, D and E). These results indicate that the two genes are likely involved in the same drug detoxification pathway in *Bcl11b*^+^ cells. This is in line with the general concept that xenobiotics can undergo chemical modification and conjugation through phase I/II drug-metabolizing enzymes (e.g., *Aldh3a1*) in order to increase their water solubility, thus facilitating their excretion through phase III transporters (e.g., *Abcb1b*) (*60, 61*). Collectively, these results suggest that *Bcl11b* controls multiple pre-existing and adaptive drug-resistance pathways, including a drug detoxification mechanism, to offer a comprehensive defense against chemotherapy.

Intriguingly, the independent CytoTRACE analysis revealed drug-treated *Bcl11b*^Hi^ cells as the least differentiated cells among the tumor subpopulations (**Fig. 4, J and K**). This indicates that the *Bcl11b*^Hi^ subset contains cells in an even more undifferentiated state in response to treatment, increasing cell fitness under pressure. This is consistent with the observed adaptive gene expression response in these cells. We also examined whether the *Bcl11b*^+^ immature basal-like cell state differs from the Epithelial-to-Mesenchymal Transition (EMT) state, which has been frequently connected to treatment resistance in cancer (*62*). The EMT process typically involves the loss of epithelial markers (*Epcam* and *Cdh1*) and the activation of EMT-specific master regulators, particularly *Snai1* and *Zeb1* (*63, 64*). Here in the tumor organoids, we found that the *Bcl11b*^+^ basal-like tumor cells highly expressed epithelial cell marker genes (*Epcam* and *Cdh1*), and were largely negative for the EMT activators, *Snai1* and *Zeb1* (fig. S9, A and B). Other EMT markers associated with a more mesenchymal phenotype, such as *Vim* and *Fn1*, were either lowly expressed in the tumor cells or displayed a non-specific expression pattern among tumor cells (fig. S9B). FACS analysis has also confirmed that the basal-like tumor cells stained positive for EpCAM (fig. S3A). Moreover, gene set enrichment analysis revealed that the normal basal cell signature was significantly enriched in the basal-like tumor cells, while the EMT gene set was expressed at comparable levels between basal and luminal cells (fig. S9C). This implies that the basal-like tumor cells exhibit more similarities to normal basal cells as opposed to cells that have undergone EMT activation. It should be noted that a small number of luminal tumor cells displayed expressions of *Snai1*, *Zeb1*, and *Zeb2* (fig. S9, A and B), which is consistent with the idea that the EMT state might arise from luminal cells at a late stage (*63*). Nevertheless, our results show that the *Bcl11b*^+^ drug-resistant tumor cells maintain epithelial cell identity and are distinct from the previously reported chemoresistant EMT states.

### The emergence of persister cells is repressed by TNF*α* treatment

Because the BCL11B^+^ cancer cells express various disparate drug-resistant programs that would require multiple therapeutics for targeting, we therefore asked whether there was a way to directly inhibit the BCL11B^+^ cell state and whether this could be a therapeutic approach against chemoresistance. To search for functional regulators of the *Bcl11b*^+^ cells, we analyzed the bulk RNA-seq data of *Bcl11b*^+^ and *Bcl11b*^-^ cells and identified TNFα signaling as the top-enriched pathway in *Bcl11b*^-^ cells (**Fig. 5A**). This finding was corroborated by the analyses of scRNA-seq data of tumor organoids and bulk RNA-seq data of WT and *Bcl11b* knockout tumor cells (**Fig. 5B**). TNFα, a pro-inflammatory factor, plays a physiological role in regulating mammary gland development (*65*) and can signal through diverse pathways including nuclear factor NF-κB and MEK/ERK pathways (*66*). Notably, single-cell analyses of normal mammary epithelium and the tumor organoid revealed an opposite expression pattern of *Tnf* and *Bcl11b* (fig. S10A). *Tnfaip2*, which is a primary response gene of TNFα, is also specifically enriched in *Bcl11b* negative cells (fig. S10A). Thus, we hypothesize that TNFα, which can be secreted by luminal cells and niche cells, could function as a natural inhibitor of the *Bcl11b*^+^ cell state.

**Fig. 5.**
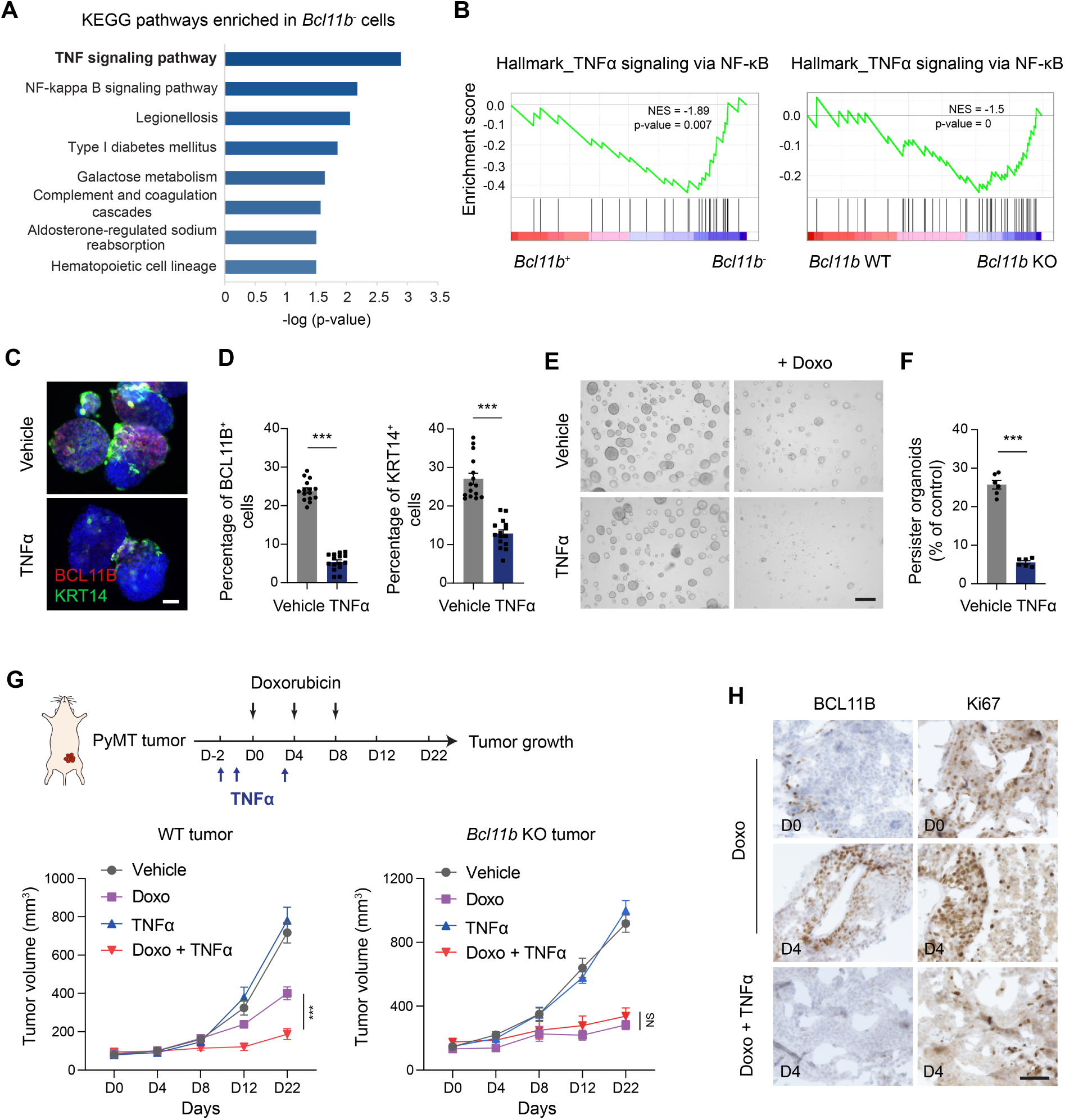
TNF*α* enhances chemotherapy efficacy by disrupting the BCL11B^+^ cancer cell state. (**A**) KEGG pathway analysis of the upregulated genes in *Bcl11b*^-^ PyMT tumor cells. (**B**) Left panel: GSEA analysis of the scRNA-seq data demonstrates increased TNFα signaling signature level in *Bcl11b*^-^ cells compared to *Bcl11b*^+^ cells. Right panel: GSEA analysis demonstrates increased TNFα signaling signature level in *Bcl11b* KO cells compared to WT cells. (**C**) Representative immunostaining images of the control and TNFα-treated tumor organoids. PyMT tumor organoids were treated with TNFα (200 ng/ml) for 6 days and analyzed for BCL11B and KRT14 expression. Scale bar, 50 μm. (**D**) Percentage of BCL11B^+^ cells or KRT14^+^ cells in tumor organoids with and without TNFα treatment; n = 15 organoids from 3 experiments. (**E**) Tumor organoids were either left untreated or pre-treated with TNFα (200 ng/ml) for 2 days before exposure to doxorubicin (200 nM). After 24-hour drug treatment, single cells from the indicated groups were replated for organoid formation under normal culture conditions. (**F**) Quantification of the frequency of drug-resistant persister organoids (% of control). n = 6 cell culture wells from 3 experiments. (**G**) Upper panel: Schematic diagram showing the *in vivo* drug treatment experimental design. NSG mice bearing PyMT tumors were treated with saline, doxorubicin (1 mg/kg, three cycles), TNFα (100 µg/kg, three doses), or doxorubicin in combination with TNFα. Lower panel: Tumor growth curves showing growth of WT or *Bcl11b* KO tumors under different treatment conditions. n = 6 mice per group, two-way ANOVA. (**H**) Representative images of BCL11B and Ki67 staining in PyMT tumors under different treatments on the indicated days. Scale bar, 50 μm. Nuclei were counter-stained with DAPI (blue). Data are presented as mean ± SEM. *p < 0.05, **p < 0.01, ***p < 0.001.

TNFα treatment caused a marked decrease in BCL11B expression in the murine mammary epithelial cell line, Comma-Dβ cells (fig. S10B). Interestingly, induced overexpression of BCL11B, in turn, dramatically attenuated TNFα-induced ERK activation in these cells (fig. S10C). This revealed an unexpected reciprocal inhibition between TNFα signaling and BCL11B in mammary epithelial cells. We then exposed PyMT tumor organoids to TNFα treatment and observed a significant reduction in the number of BCL11B^+^ tumor cells within a few days (**Fig. 5, C and D**). The percentage of KRT14^+^ cells also decreased accordingly in the post-treatment tumor organoids (**Fig. 5, C and D**). We next treated single *Bcl11b*^tdTomato^-positive tumor cells with TNFα and tracked the fates of these single cells. In the absence of TNFα, single BCL11B^+^ cells developed into tumor organoids with a prominent population of KRT14^+^ basal-like cells among the luminal cells (fig. S10D). Treatment with TNFα did not compromise the organoid formation efficiency, but notably decreased the proportion of KRT14^+^/ BCL11B^+^ basal-like cells within the formed organoids (fig. S10, D and E). This indicates a TNFα-induced basal to luminal lineage shift in the tumor cells. Importantly, pre-treatment with TNFα effectively decreased the emergence of drug-resistant persister cells (**Fig. 5, E and F**), supporting the idea that impairing the BCL11B^+^ cell state can avert the development of drug resistance in breast cancer.

To assess the therapeutic potential of the TNFα combination treatment *in vivo*, we administered three doses of TNFα in conjunction with three cycles of doxorubicin to tumor-bearing recipient mice (**Fig. 5G**). TNFα injection was given at a tolerated dose based on mouse studies and human clinical trials (*67, 68*) and was started prior to the chemotherapy treatment in order to eradicate BCL11B^+^ tumor cells before chemotherapy was administered. TNFα administration alone did not induce noticeable alterations in tumor growth or the overall health of the mice. However, when combined with doxorubicin, TNFα remarkably enhanced the efficacy of chemotherapy in suppressing the growth of the WT tumors (**Fig. 5G**). The combination of TNFα and doxorubicin had no synergistic effects on *Bcl11b*-deficient tumor cells, suggesting that TNFα mainly functioned through targeting the BCL11B^+^ cells (**Fig. 5G**). Analysis of the on-treatment and post-treatment samples revealed that doxorubicin exposure caused a significant decrease in cell proliferation in the tumor, although residual zones of Ki67^+^ cells were consistently observed during the treatment (**Fig. 5H** and fig. S10F). BCL11B^+^ cells were also found to be enriched in these residual lesions (**Fig. 5H** and fig. S10G). In contrast, treatment with a combination of TNFα and doxorubicin eliminated both BCL11B^+^ cells and residual proliferating cells (**Fig. 5H** and fig. S10, F and G). Together, these results reveal that pre-treatment with TNFα prior to chemotherapy can strongly enhance chemotherapy efficacy by impairing the BCL11B^+^ cell state and preventing the emergence of cancer persisters.

### BCL11B promotes treatment resistance in human breast cancer

To assess the implications of our mouse tumor findings for human breast cancer, we first revisited the single-cell breast cancer atlas. Considering the potential dropout of *BCL11B* expression in single-cell datasets due to its low expression levels in cells, we broadly divided tumor cells into *BCL11B*^High^ and *BCL11B*^Low^ compartments based on the average expression level of *BCL11B* in each Seurat cell cluster (fig. S11A). As expected, *BCL11B* expression was enriched in the *BCL11B*^High^ compartment (**Fig. 6A**). We next defined a ‘BCL11B signature’ that consists of the top 10 BCL11B-associated genes (*Ephb1*, *Dsg3*, *Col17a1*, *Tgfbi*, *Cntfr*, *Ngfr*, *Cyp26a1*, *Postn*, *Pnpla1*, and *Id4*) from the mouse tumor data. Genes that were highly expressed in *Bcl11b*^+^ cells and were markedly downregulated by *Bcl11b* deletion were identified as BCL11B-associated genes (fig. S11B). We then validated the BCL11B signature in our scRNA-seq dataset of the tumor organoids and found a strong correlation between *Bcl11b* expression and the BCL11B signature (fig. S11C). Upon examining the cancer epithelia in the breast cancer atlas, we observed that the BCL11B signature and *KRT14* were significantly enriched in the *BCL11B*^High^ compartment, while *KRT18* was expressed at a lower level in the *BCL11B*^High^ cells (**Fig. 6A**). The genes associated with the rudimentary basal-like cell state and the drug resistance programs, including *DSG3*, *TGFBI*, and *ALDH3A1*, were also enriched in the *BCL11B*^High^ tumor population (fig. S11D). Moreover, the *BCL11B*^High^ tumor compartment showed no enrichment in myoepithelial markers and displayed a lower level of TNFα signaling activity (**Fig. 6A** and fig. S11E). Together, these results suggest that the TNFα/BCL11B^+^ immature cell state/drug-resistance axis is largely conserved in human breast cancer. We also observed that the majority of cancer cells were positive for *EPCAM* (**Fig. 6A** and fig. S11E) and demonstrated low levels of *SNAI1* and *ZEB1*, two critical EMT factors in breast cancer (fig. S11E), consistent with the findings in mouse tumors.

**Fig. 6.**
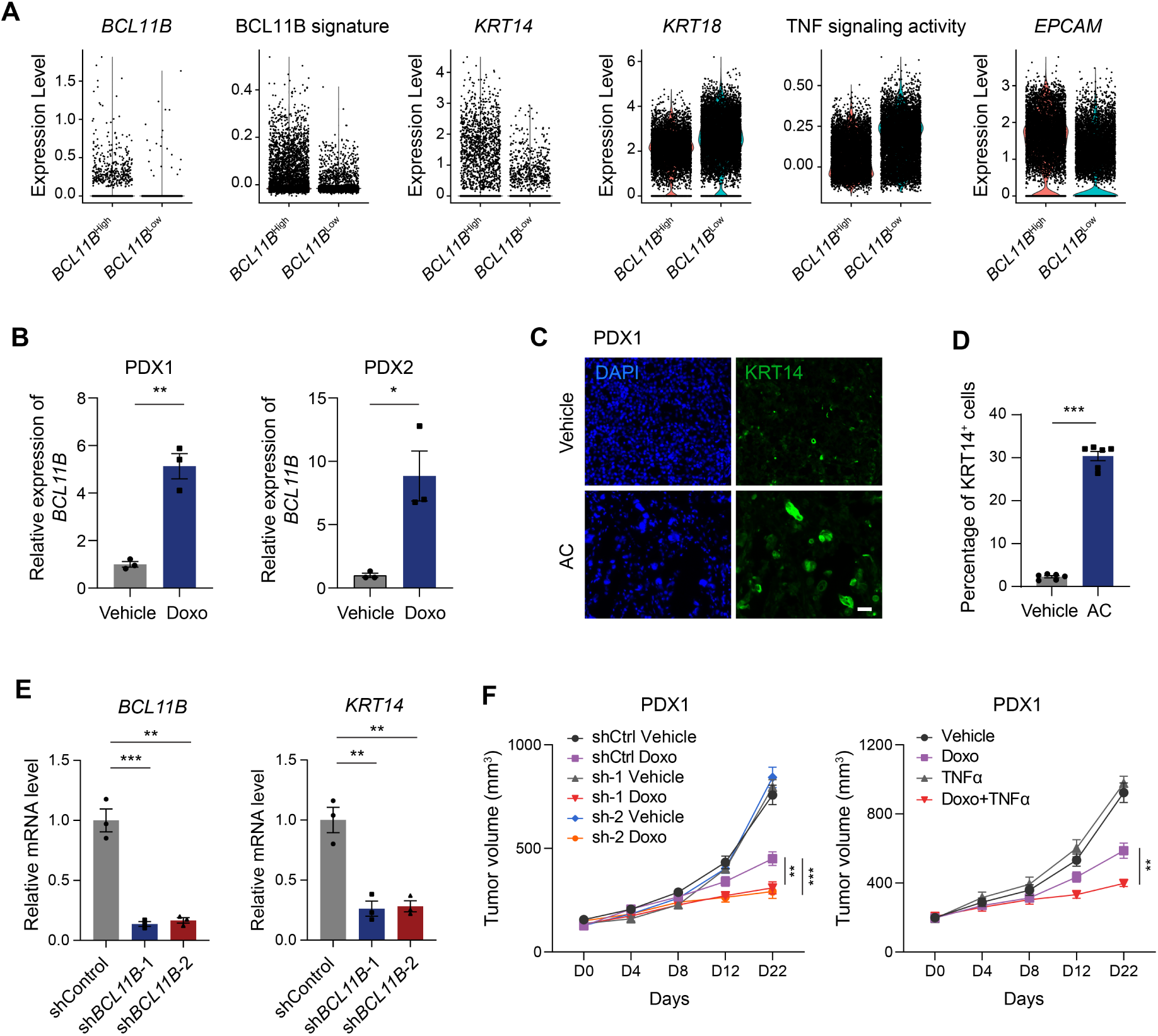
BCL11B promotes chemoresistance in human breast cancer. (**A**) Violin plots comparing the expression levels of *BCL11B*, BCL11B signature, *KRT14*, *KRT18*, TNF signaling activity, and *EPCAM* between the *BCL11B*-high epithelial compartment and *BCL11B*-low epithelial compartment in the single-cell breast cancer atlas. (**B**) Real-time PCR quantification of *BCL11B* mRNA levels in cancer cells isolated from control PDXs and doxorubicin-treated PDXs. *GAPDH* was used as an internal control. Doxorubicin was administered at three doses (once every 4 days) and tumor cells were collected 1 day after the final dose. n = 3 mice per group. (**C**) Representative images of KRT14 immunostaining in control and drug-treated PDX1 samples. Scale bar, 50 μm. (**D**) Percentage of KRT14^+^ cells under the indicated conditions; n = 3 tumors. (**E**) Real-time PCR quantification of *BCL11B* and *KRT14* mRNA levels in control PDX1 and *BCL11B* knockdown PDX1 cells. n = 3 experiments. (**F**) Left panel: tumor growth curves showing the growth of WT PDX1 tumors or *BCL11B* knockdown PDX1 tumors upon saline (Vehicle) and doxorubicin (Doxo) treatment. Right panel: tumor growth curves showing the growth of PDX1 tumors upon saline (Vehicle), doxorubicin (Doxo), TNFα, or doxorubicin in combination with TNFα treatments. Treatment schemes are the same as those described in Figure 5G. n = 6 mice per group, two-way ANOVA. Nuclei were counter-stained with DAPI (blue). Data are presented as mean ± SEM. *p < 0.05, **p < 0.01, ***p < 0.001.

To directly assess the functional role of BCL11B in promoting drug resistance in human breast cancer, we further utilized the breast cancer patient-derived xenograft (PDX) models that were previously established in our laboratory (*69*). High levels of BCL11B^+^ tumor cells were detected in 2 out of the 5 triple-negative xenograft models we tested, consistent with our findings that BCL11B is expressed in a subset of human breast cancers. Chemotherapeutic treatment induced upregulation of *BCL11B* mRNA levels in both PDX tumor lines, which was corroborated by the observation that KRT14^+^ cells were significantly enriched in the post-treatment tumors (**Fig. 6, B-D**). We then knocked down *BCL11B* in the PDX models and found a corresponding decrease in the expression of *KRT14* (**Fig. 6E** and fig. S12A). Similar to the findings in mouse studies, the anti-tumor efficacy of doxorubicin was significantly augmented upon the inhibition of BCL11B through shRNA-mediated knockdown in the PDX models (**Fig. 6F** and fig. S12B). We also confirmed that TNFα treatment notably enhanced the anti-tumor effects of chemotherapy on WT PDX tumor cells (**Fig. 6F**). We next extended our analysis to human breast cell lines. Analysis of the breast cancer cell lines in the DepMap (*70*) database showed that *BCL11B* displayed significant variability in expression levels, with high expression in a subgroup of the cell lines (fig. S12C), including BT20, HCC70, and HCC1395. Interestingly, almost all of these *BCL11B*^High^ cancer cell lines demonstrated a notable resistance to commonly used chemotherapeutic drugs, as revealed in the GDSC database (*71*) (fig. S12C). To verify whether BCL11B can initiate the drug-resistance programs in the human context, we induced an overexpression of BCL11B in MCF10A normal human breast cells. These cells have minimal gene mutations and serve as a model for both normal and malignant mammary epithelial cell biology (*72*). The *ex vivo* drug treatment assay showed that the ectopic expression of BCL11B induced *de novo* emergence of persister organoids in response to doxorubicin treatment (fig. S12, D and E). Overall, our studies on patient-derived models revealed that BCL11B^+^ cancer cells functionally promote resistance against chemotherapy in human breast cancer.

## Discussion

Different mechanisms mediating chemotherapy resistance in cancer have been proposed, but a key unresolved question is why tumors display great variability in drug responses. A well-appreciated theory of drug-tolerant persister proposes that a subpopulation of tumor cells can enter a slow-cycling, drug-tolerant state under drug pressure to evade treatment (*3*). It has been suggested that cancer persisters could emerge non-selectively from tumor cells (*73, 74*), but it remains challenging to explain the reasons behind the significant variation in tumor responses to chemotherapy. Evidence from recent lineage-tracing studies in colorectal cancer and skin cancer reveals that certain pre-existing stem-like tumor subpopulations persist during chemotherapy and drive tumor regeneration (*9–11*), supporting the notion that the emergence of cancer persisters can be driven by pre-existing mechanisms, especially in highly heterogeneous cancers.

Here in breast cancer, we revealed that the pre-existing BCL11B^+^ cancer cells can selectively transition to a persister state following treatment, benefiting from their fitness advantages conferred by their innate drug metabolism machinery. Interestingly, the BCL11B^+^ cancer cells also exhibit a notable capacity to adapt transcriptionally in response to drug treatment, whereas the luminal tumor cells are prone to entering cell cycle arrest and apoptosis under drug stress. Therefore, our results support a model in which pre-existing tumor populations with inherent fitness advantages can preferentially and actively evolve a resistant state through transcriptional adaptation mechanisms (**Fig. 7**). This is distinct from the general notion that drug resistance emerges through the passive selection of pre-existing subpopulations or through transcriptional adaptations that occur stochastically among tumor cells (*2*). Notably, we have also shown that tumors with populations of BCL11B^+^ cells first decrease in size after therapy, but the minority population of BCL11B^+^ cells will regenerate a tumor with a mixed population of cells, which will once again shrink after the same therapy (**Fig. 2L**, and fig. S3, G and H, and fig. S6A). This provides an alternative model explaining why some patients initially respond to therapy but their tumors continue to regrow after a drug holiday and exhibit shrinkage again upon the reintroduction of the same therapy (*75*).

**Fig. 7.**
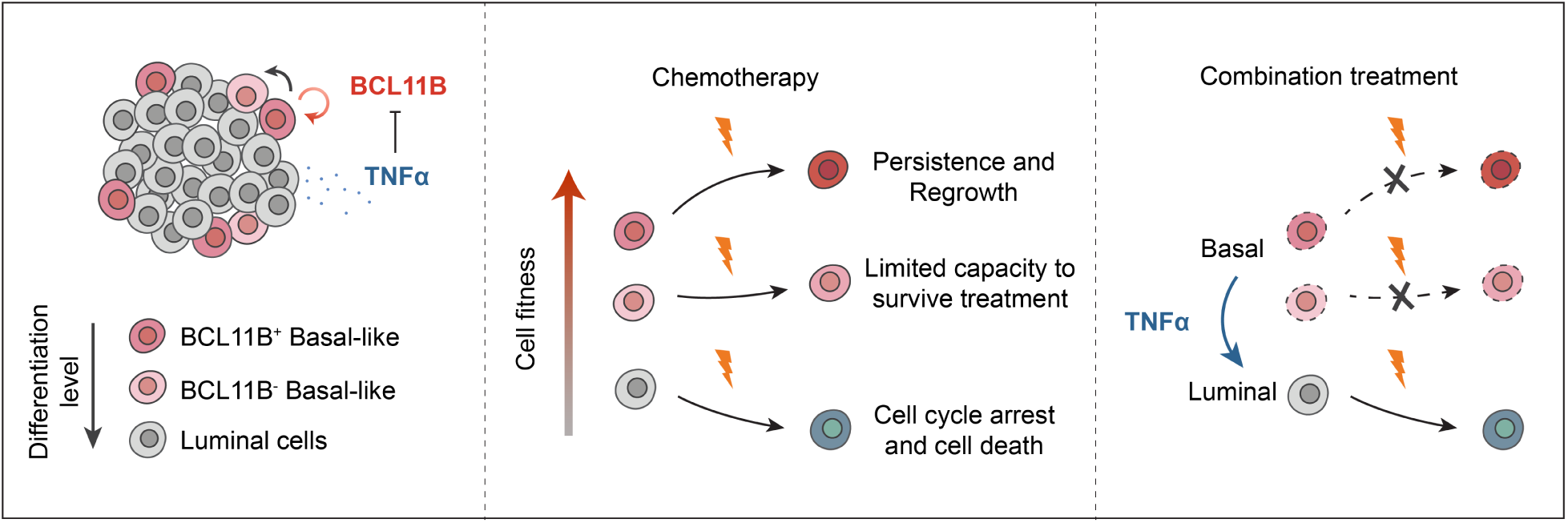
BCL11B predetermines a drug-resistant persister state in breast cancer that is reversed by TNF*α* treatment. Schematic illustration depicting that BCL11B functionally maintains an immature basal-like tumor population, with BCL11B^+^ cells being in the more immature state and possessing a superior capability to transition to a drug-resistant persister state. The BCL11B^-^ basal-like tumor cells retain some level of drug resistance, while luminal tumor cells are drug sensitive and readily undergo cell cycle arrest and cell death upon treatment. TNFα can inhibit BCL11B expression and drive the cells out of the drug-resistant basal-like state, thus significantly improving chemotherapy efficacy.

As a zinc finger transcriptional factor and a component of various chromatin-remodeling complexes such as NURD and SWI/SNF (*21, 76*), BCL11B regulates different gene programs depending on the cellular context and cell type. BCL11B is essential for the fetal development of multiple systems and also plays a crucial role in maintaining stem cells and progenitors in certain adult tissues including the mammary gland, epidermis, and neural tissues (*17–19*). Consistent with its context-dependent role, BCL11B has been found to be overexpressed and exhibit oncogenic functions in Ewing sarcoma and head and neck squamous cell carcinoma, whereas in T-cell Acute Lymphoblastic Leukemia (T-ALL), BCL11B is recognized as both a haploinsufficient tumor suppressor and an oncogene (*22, 23, 77*). The recent reported role of BCL11B in base excision repair in T cells offers a possible explanation for its paradoxical roles in T-ALL (*78*). BCL11B can prevent the acquisition of oncogenic mutations in normal cells, but it can also protect cancer cells from oxidative DNA damage. This aligns with the observation that both the loss-of-function mutations in BCL11B (18%) (*79*) and the protein overexpression are prevalent in T-ALL. In contrast, loss-of-function mutations in BCL11B are rare in many other malignancies including human breast cancers (less than 1%). Here in breast cancer, we identified a previously unrecognized role of BCL11B in pre-determining a drug-resistant persister state and exposed new mechanisms through which BCL11B safeguards cancer cells from chemotherapy treatment, thus revealing a promising target in combating chemoresistance. The potential role of BCL11B in protecting breast cancer cells from DNA damage, including by regulating ROS through GSH metabolism, is also an intriguing direction for future investigation.

Breast cancer is a disease that is widely recognized for its significant level of heterogeneity. The identification of biomarkers such as HER2/NEU, which can pinpoint a specific subset of patients who can benefit from targeted therapy, can be critical for achieving successful treatment outcomes (*33*). A number of other marker genes that are enriched in less differentiated breast cancer cells have been identified, but many of them lack specificity, limiting their predictive value and potential for therapeutic targeting (*80*). Here, by systematically interrogating single-cell atlases of human breast cancer using the CytoTRACE algorithm, we identified a previously unrecognized *BCL11B*^+^ immature cancer cell state that is naturally impervious to a multitude of chemotherapeutic drugs, resulting in a higher possibility of chemotherapy resistance. Therefore, our findings identify *BCL11B* as a new candidate biomarker for subtyping breast cancer and predicting chemotherapy resistance in some patients. Utilizing BCL11B as an immunohistochemical prognostic marker for assessing biopsy samples prior to the start of chemotherapy shows great promise and feasibility, although it will necessitate prospective clinical studies for validation.

It is intriguing how multiple mechanisms can converge within a single cancer cell population to confer drug resistance. Notably, directly targeting this master regulator of the drug-resistant cell state through a therapeutic agent, TNFα, could eliminate the emergence of drug resistance and greatly enhance chemotherapy efficacy in the subset of breast cancer tumors with BCL11B^+^ cancer cells (**Fig. 7**). In line with this, early research has illustrated the success of utilizing TNFα in conjunction with chemotherapy for the treatment of advanced soft tissue sarcomas (*68, 81*). Treatment of breast cancer patients, particularly TNBC patients with cancers localized to the breast and regional lymph nodes with preoperative chemotherapy including taxanes and doxorubicin results in increased survival, especially in patients whose breast tumors demonstrate a major response to therapy (*82*). Our results show that in tumors with higher levels of BCL11B^+^ cancer cells, adding TNFα prior to the start of chemotherapy treatment could dramatically improve treatment response. Thus, BCL11B might hold promise as a biomarker for identifying a subset of human breast cancers with a higher risk of developing chemoresistance that, like some patients with advanced soft tissue sarcomas, may benefit from precision therapy combining chemotherapy with TNFα.

TNFα has failed in some clinical trials because it does not show efficacy as a single agent and high doses have significant toxicity. Our data identifies a potential way to overcome these issues. We show that treatment with TNFα alone is not effective in preclinical models, consistent with Phase 1 clinical trial data. However, pretreatment with low doses of TNFα, which have not shown toxicity in clinical trials (*68*), sensitizes tumors to chemotherapy. TNFα administration concurrent with chemotherapy would not show efficacy because tumors would not have downregulated BCL11B expression and chemotherapy resistance. Based on our data, there is also great promise in future efforts aimed at developing new TNFα agents that can selectively target cancer cells to minimize potential systemic side effects. We have also found that targeting the drug detoxification program with small molecule inhibitors shows promise in reversing the emergence of drug-resistant organoids, but we were unable to test it *in vivo* due to considerable toxicity (data not shown). Developing specific inhibitors with reduced toxicity could be a promising direction for future work, but their translational values may also be restricted due to the potential compensation from other drug-resistance mechanisms in BCL11B^+^ cells.

Cancer treatments like conventional chemotherapy not only affect tumor cells but also have profound impacts on the tumor microenvironment. Our studies with organoid cultures and xenograft models do not fully recapitulate the complex drug responses in human cancers. Further studies are warranted to understand the interplay between the tumor microenvironment and BCL11B^+^ cells during drug treatment. We acknowledge that there might be other mechanisms contributing to the development of chemoresistance in breast cancers that have low BCL11B expression, but our results highlight a critical role of BCL11B^+^ cells in predisposing cancer cells to evade treatment in at least a subset of breast tumors, thus revealing a promising therapeutic target.

## Materials and Methods

### Mice

MMTV-PyMT (stock # 002374 and 022974) and NOD *scid* gamma (NSG, stock # 005557) mice were purchased from Jackson Laboratories. *Bcl11b*^tdTomato^ knock-in reporter mice were generously provided by Pentao Liu’s lab. For xenograft studies, 8-12 weeks old immunocompromised NSG females were used. Mice were maintained in-house under aseptic sterile conditions. Mice were administered with autoclaved food and water. All animal procedures were conducted in accordance with a protocol (10868) approved by the Stanford University APLAC committee.

### Cell lines

MCF10A and 293T cells were obtained from ATCC. MCF10A cells were grown in DMEM (Invitrogen) supplemented with 5% Horse Serum (Invitrogen), 20 ng/ml EGF (Invitrogen), 0.5 mg/ml Hydrocortisone (Sigma), 100 ng/ml Cholera Toxin (Sigma), 10 µg/ml Insulin (Sigma), and 1% PSA (Invitrogen). 293T cells were grown in DMEM (Invitrogen) supplemented with 10% FBS (Hyclone), Glutamax (ThermoFisher Scientific), sodium pyruvate (Life Technologies), and 1% PSA (Invitrogen). Comma D beta cell line was cultured in DMEM-F12 (Invitrogen) supplemented with 2% Fetal Bovine Serum (Hyclone), 1% PSA (Invitrogen), 10 ng/ml EGF (Invitrogen) and 5 µg/ml Insulin (Sigma). All cells were incubated at 5% CO_2_ and 37°C. None of the cell lines used is listed in the database of commonly misidentified cell lines maintained by ICLAC. Cell lines have not been authenticated but all cell lines used were passaged less than 10 times from when the original cells from the vendors were thawed.

### Deconvolution of bulk breast tumors and survival analysis

CIBERSORTx was used to deconvolve cell type abundances from microarray gene expression data of bulk breast tumors from the METABRIC dataset and Hatzis dataset (n = 508). Default parameters as described in the “Tutorial” page at http://cibersortx.stanford.edu/ were used to generate a signature matrix from the scRNA-seq dataset of breast cancers (*26*). Quantile normalization was run on microarray data, and bulk-mode batch correction (B-mode) was applied for cross-platform deconvolution. The R package survival (v3.1.12) was applied to determine the correlation of indicated cancer populations with relapse-free survival. The samples were stratified into two groups, “high” and “low,” based on whether the abundance of the deconvolved cancer cell population was more or less than the median. A Cox proportional hazard model was then employed to calculate the effect of the abundance of the indicated cancer cell populations on Relapse-free survival or Distant recurrence-free survival. Patients were censored at 60 months for the survival analysis.

### Preparation of single-cell suspension and flow cytometry

Mouse primary mammary tumors were obtained from 12-16 weeks-old virgin female MMTV-PyMT mice. Mouse or human xenograft samples were obtained from NSG recipient mice that had been transplanted with tumor cells. For human samples, informed consent was obtained, and human normal breast and breast tumor specimens were collected according to guidelines from Stanford University’s Institutional Review Board (4344). Tumor samples were dissected and processed according to the published protocol (*69*) with minor modifications. In brief, samples were mechanically dissociated into small pieces (1-2 mm^3^) with a razor blade and then digested at 37°C with 1,500 U collagenase and 500 U hyaluronidase (Stem Cell Technology) in Advanced DMEM/F-12 (Thermo Fisher Scientific) supplemented with 2% FBS (Hyclone), and 1% PSA (Invitrogen) for 3 hours with hourly pipetting. After digestion, the reaction was neutralized by adding FACS staining buffer (HBSS + 2% FBS + 1% PSA) and spun down at 1500 rpm for 5 min. The pellet was then treated with 5 mL of ACK lysis buffer (Lonza) on ice for 5 min to deplete red blood cells. The sample was spun down and the cells were further digested in 4ml 0.25% Trypsin-EDTA (GIBCO) supplemented with 1,000 U DNase for 10 min at 37°C. Dissociated cells were filtered by a 40 μm strainer and washed with FACS staining buffer. For FACS analysis and sorting, single cells were stained with CD45 (Biolegend), CD31 (Biolegend), Ter119 (Biolegend), CD49f (Biolegend), EpCAM (Biolegend), and CD24 (Biolegend) with appropriate conjugated fluorophores for 20 min on ice. The cells were then washed and resuspended in FACS staining buffer containing DAPI (1 µg/ml). Stained cells were analyzed and sorted on a FACSAria II (BD Bioscience) instrument with a 100 μm nozzle. A complete list of antibodies is provided in table S5.

### Organoid culture

20ul of growth factor reduced Matrigel (BD Bioscience) was added to a well of a round-bottom 96-well plate and allowed to solidify at 37°C for 10min. FACS-sorted tumor cells were then resuspended in 150 µl organoid culture media (Advanced DMEM/F12 + 2% FBS + PSA + N2 + B27 + 10 mM HEPES + EGF (10 ng/ml, Invitrogen) + Rspo1 (250 ng/ml, R&D) + ROCK inhibitor Y27632 (10 µM, Sigma) + Noggin (100 ng/ml, R&D)), and overlaid on top of the Matrigel. Organoids were grown in humidified tissue culture incubators at 37°C in 5% CO_2_ and were supplemented every two or three days with fresh media. For passaging, organoids were first incubated in pre-chilled cell recovery solution (Corning) on ice for 20 min and then dissociated into single cells by trypsin digestion.

For lentivirus-mediated gene knockdown, organoid cells were transduced with scrambled virus (Sigma), *Aldh3a1*, or *Abcb1b* shRNA virus that was produced in 293T cells as described previously (*83*). Single organoid cells were mixed with concentrated lentiviral supernatant (to a MOI. of 20) in organoid culture media with 4 µg/ml of Polybrene (Sigma-Aldrich) and incubated in a 48-well non-adherent culture plate overnight. The next day, cells were washed and cultured on Matrigel in normal organoid media. 72 hours later, positive cell selection was started by puromycin addition.

### CRISPR-mediated genome editing in tumor organoids

Guide RNA targeting the exon 1 of *Bcl11b* (CCGGGCAATGTCCCGCCGCAAACAGG) was cloned into Cas9 containing lentiCRISPR v2 vector. Lentivirus production and transfection were performed as described above. After infection, single tumor cells were plated on Matrigel for organoid culture, followed by selection with puromycin. Single organoid clones were then isolated and expanded. Genomic DNA was isolated from the selected clone and the targeted mutation in the *Bcl11b* locus was confirmed by Sanger sequencing. Two single organoid clones were verified through immunostaining and subsequently utilized for the experiments.

### Transplantation

Dissociated single cells from tumors were resuspended in 100 µl of media (organoid culture media + 50% of Matrigel) and injected into the nipple of female NSG mice at the fourth mammary fat pad. For lentivirus-mediated *BCL11B* knockdown transplant assay, dissociated xenograft cells were transduced overnight in organoid culture media with 20 MOI. of lentivirus. The next day, PDX cells were transferred to Matrigel and cultured in organoid media + Neuregulin 1 (5 nM, PeproTech) + A83-01 (500 nM, Tocris) + SB202190 (500 nM, Sigma). 72 hours after transduction, organoids were selected with 1 µg/mL puromycin. Transduced organoid cells were grown in the selection media for 4-5 days and then transplanted to NSG recipient mice as described above. A summary of the PDX models used in this study is described in table S6.

### *In vivo* treatment

Tumor-bearing NSG mice were randomized into control and treatment groups when the tumor width reached 3-5 mm for drug treatment. Saline, doxorubicin (1 mg/kg body weight, every 4 or 6 days), or doxorubicin (1 mg/kg) combined with cyclophosphamide (60 mg/kg) were given to the mice by intraperitoneal injection for the duration indicated in the regimen. For experiments involving TNFα treatment, murine or human recombinant TNFα (100 µg/kg, PeproTech) was administered by intraperitoneal injection as indicated in the regimen. Health and body weight were monitored every 2-3 days over the course of treatment, and tumor growth was measured every 3-4 days until the endpoint of the experiment.

### RNA isolation and real-time PCR

Total RNA was extracted with RNeasy Plus Micro Kit (Qiagen) and reverse transcribed to cDNA using SuperScript III First Strand Synthesis kit (Life Technologies) according to the manufacturer’s instructions. Real-time PCR was performed on the 7900HT Fast Real-Time PCR System (Applied Biosystems) or the QuantStudio 7 PRO Real-Time PCR System (Thermo Fisher) using Power SYBR Green PCR Master Mix (Life Science Technologies). *Gapdh* or *Actb* was used to normalize the expression values. PCR primers are listed in table S7.

### RNA-sequencing and data processing

20-100 ng total RNA was used to generate barcoded RNA-seq libraries using the NEBNext Single Cell/Low Input RNA Library Prep Kit for Illumina (New England Biolabs) following the manufacturer’s instructions. Libraries were sequenced on a NextSeq 550 (Illumina) or HiSeq 2500 (Illumina) to generate 75bp SE reads or 76bp PE reads. The quality of trimmed Fastq reads was assessed using FastQC. HISAT2 was then used to align the RNA-seq reads to the mouse reference genome (mm10), followed by gene count quantification using htseq-count (*84, 85*). Alternatively, RNA-seq results were aligned to the mouse reference genome (mm10) and quantified using Salmon (*86*). Count matrices were analyzed with DESeq2 (*87*) or Degust (http://degust.erc.monash.edu). Heatmaps were generated using Morpheus software (Broad Institute). KEGG pathway analysis was performed using EnrichR (http://amp.pharm.mssm.edu/Enrichr/).

### Single-cell RNA sequencing

Pre-grown tumor organoids were treated with or without doxorubicin (100 nM) for 24 hours. After the treatment, tumor organoids were recovered from Matrigel and digested into single cells using 0.25% Trypsin-EDTA (GIBCO) supplemented with DNase I for 15 min at 37°C. Freshly sorted live single cells were counted and loaded onto the Chromium Controller (10x Genomics). Each sample was loaded into a separate channel on the Single Cell 3′ Chip targeting 3,000 cells. Libraries were prepared using the Chromium Single Cell 3′ Reagent kits v3 (10x Genomics) following the manufacturer’s recommendations. The multiplexed libraries were loaded on a NovaSeq 6000 (Illumina) and paired-end reads were produced using a 200-cycle kit. Each sample was sequenced to an average depth of ∼40,000 read pairs per cell.

### Single-cell RNA-sequencing data bioinformatic analysis

The Cell Ranger pipeline was used to process the raw sequencing data, producing count matrices initially containing 3,509 cells in the Control group and 3,576 cells in the Doxo group. Package Seurat v4 was used in R to perform downstream analyses (*88*). Gene signature scores for the “Basal signature”, “Luminal signature”, “Myoepithelial signature” and “TNF signaling activity” were calculated using the AddModuleScore function in Seurat. The “TNF signaling activity” is composed of the TNF signaling genes in the HALLMARK_TNFA_SIGNALING_VIA_NFKB gene set. Single-cell level prediction of differentiation states in tumor organoid cells was performed in R using the CytoTRACE package publicly available at https://cytotrace.stanford.edu. Single-sample enrichment of the “Normal Basal Signature,” “Normal Luminal Signature” and the dbEMT (*89*) gene sets were analyzed using ssGSEA as implemented in the escape package (*90*). Signature genes (top 200 DE genes for each lineage) for basal and luminal cells from normal mammary glands were obtained from publicly available RNA-seq data (*91*) as described in the previous study (*92*).

### Single-cell gene expression analysis

Single-cell gene-expression analysis of normal human breast was done using Fluidigm’s M48 quantitative PCR (qPCR) DynamicArray microfluidic chips (Fluidigm) as previously described (*19, 93*).

### Immunofluorescence

Normal mammary tissues or tumor samples were fixed in 4% formaldehyde solution and subjected to either paraffin or O.C.T. embedding (Tissue-Tek). Paraffin sections were de-paraffinized, dehydrated, and microwaved for 15 min at 95 °C in Sodium Citrate Buffer (10mM Sodium Citrate, 0.05% Tween 20, pH 6.0) for antigen retrieval. Frozen sections were first hydrated with PBS for 10min at room temperature and then microwaved for 15 min at 95 °C in Sodium Citrate Buffer for antigen retrieval. Tissue sections were blocked with staining buffer (PBS + 5% BSA + 0.1% Triton X-100) for 1 hour at room temperature and then incubated overnight at 4 °C with the primary antibodies diluted in the staining buffer. Samples were subsequently washed with PBS and incubated with conjugated secondary antibodies (Invitrogen) at 1:300 in staining buffer + DAPI (1 µg/ml) for 1 hour at room temperature (antibodies are listed in table S5). After PBS washing, tissue sections were mounted with Fluoromount® Aqueous Mounting Medium (Sigma). Images were acquired by a Carl Zeiss LSM 710 Meta confocal microscope or a Keyence fluorescence microscope.

For whole-mount organoid staining, pre-chilled cell recovery solution (Corning) was added to the well for 20 min on ice to retrieve the organoids. The organoids were then transferred to a microcentrifuge tube and pelleted by quick spinning using a mini centrifuge. After being fixed in 4% formaldehyde solution for 20 min, the recovered organoids were permeabilized and blocked in organoid staining buffer (PBS + 5% BSA + 1% Triton X-100) for 1 hour at room temperature, followed by overnight incubation with primary antibodies diluted in the organoid staining buffer at 4 °C. The organoids were then washed with PBS and incubated with conjugated secondary antibodies (Invitrogen) at 1:300 in organoid staining buffer + DAPI (1 µg/ml) for 1 hour at room temperature. After PBS washing, the organoids were mounted on a glass slide for immunofluorescence imaging.

### Western blot

Cells were lysed in ice-cold RIPA buffer containing 1x Halt Protease and Phosphatase inhibitor cocktails (ThermoFisher Scientific) and boiled on a heat block at 100-degree for 10 min. Samples were loaded on a 4%-20% gradient SDS-PAGE gel (Bio-Rad) and run at 100V for 90 min and transferred onto PVDF membranes. Membranes were blocked with TBST + 5% Non-fat dry milk for 1 hour at room temperature and then incubated with primary antibodies at 4°C overnight. Membranes were subsequently washed with TBST and stained with HRP-conjugated secondary antibodies. Membranes were then washed and visualized using the SuperSignal™ West Dura Extended Duration Substrate (ThermoFisher Scientific).

### Data and materials availability

All data generated or analyzed during this study are included in this published article (and its supplementary files). The single-cell and bulk RNA-sequencing data generated in this study have been deposited in the Gene Expression Omnibus with the primary accession code GSE228442. Previously published scRNA-seq data that were reanalyzed here are available at GEO under accession codes GSE109774, GSE176078 and at the European Genome-phenome Archive (EGA) under study number. EGAS00001004809. Previously microarray expression data reanalyzed here are available at GEO under accession codes GSE6883. All bioinformatics tools used in this study are publicly available. Patient-derived xenografts used in the manuscript will be available upon request with an appropriate Material Transfer Agreement (MTA) with Stanford University.

### Quantification and statistical analysis

Data values are represented as mean ± standard error (SEM). P-values are calculated using t-test, Wilcoxon rank sum test or 2-way ANOVA test. Survival was measured using the Kaplan-Meier method. Each experiment has been replicated at least 3 times. For animal studies, the sample size was not predetermined to ensure adequate power to detect a pre-specified effect size. No animals were excluded from analyses, experiments were not randomized and investigators were not blinded to allocation during experiments.

## Author Contributions

Z.Q., S.C., and M.F.C. conceived and designed the study. Z.Q. and S.C. performed experiments and analyzed data with supervision from M.F.C. G.S.G. assisted with bioinformatic analysis. A.H.K., S.S.S., and D.Q. provided technical support. F.M.D. assisted with the collection of patient specimens. A.M.N provided guidance on the project. Z.Q. and M.F.C. wrote the manuscript. A.M.N., S.C., A.H.K., S.S.S., and W.H.D.H. assisted with manuscript review and editing.

## Acknowledgments

We thank Catherine Carswell Crumpton, Cheng Pan, and other flow cytometry staff for their help and the animal facility core members. The BD Aria instruments were funded by NHI grant S10-1S10RR02933801. We thank Margaret Cuadro for administrative assistance. We thank the Stanford Neuroscience Microscopy Service, supported by NIH 510 grant NS069375. This work was supported by the U.S. Department of Defense (W81XWH-13-1-0281 and W81XWH-11-1-0287 to MFC), the Virginia and D.K. Ludwig Fund for Cancer Research (to MFC), the Breast Cancer Research Foundation (BCRF-23-027 to MFC), the Stanford Bio-X Interdisciplinary Initiatives Seed Grants Program (IIP) (AMN, MFC), the Stanford School of Medicine Dean’s Fellowship (to ZQ).

## List of Supplementary Materials

Figs. S1 to S12

Tables S1-7 (Excel files)

## References

1. N. Vasan, J. Baselga, D. M. Hyman, A view on drug resistance in cancer. Nature 575, 299–309 (2019).

2. J. C. Marine, S. J. Dawson, M. A. Dawson, Non-genetic mechanisms of therapeutic resistance in cancer. Nat Rev Cancer 20, 743–756 (2020).

3. M. J. Hangauer et al., Drug-tolerant persister cancer cells are vulnerable to GPX4 inhibition. Nature 551, 247–250 (2017).

4. G. De Conti, M. H. Dias, R. Bernards, Fighting Drug Resistance through the Targeting of Drug-Tolerant Persister Cells. Cancers 13, (2021).

5. W. A. Weber, Assessing tumor response to therapy. J Nucl Med 50 Suppl 1, 1S–10S (2009).

6. C. Liedtke et al., Response to neoadjuvant therapy and long-term survival in patients with triple-negative breast cancer. J Clin Oncol 26, 1275–1281 (2008).

7. P. R. Prasetyanti, J. P. Medema, Intra-tumor heterogeneity from a cancer stem cell perspective. Mol Cancer 16, 41 (2017).

8. A. Skibinski, C. Kuperwasser, The origin of breast tumor heterogeneity. Oncogene 34, 5309–5316 (2015).

9. A. Alvarez-Varela et al., Mex3a marks drug-tolerant persister colorectal cancer cells that mediate relapse after chemotherapy. Nat Cancer, (2022).

10. Y. Ohta et al., Cell-matrix interface regulates dormancy in human colon cancer stem cells. Nature 608, 784–794 (2022).

11. N. Oshimori, D. Oristian, E. Fuchs, TGF-beta promotes heterogeneity and drug resistance in squamous cell carcinoma. Cell 160, 963–976 (2015).

12. K. Polyak, Heterogeneity in breast cancer. J Clin Invest 121, 3786–3788 (2011).

13. K. M. Turner, S. K. Yeo, T. M. Holm, E. Shaughnessy, J. L. Guan, Heterogeneity within molecular subtypes of breast cancer. Am J Physiol Cell Physiol 321, C343–C354 (2021).

14. N. Y. Fu, E. Nolan, G. J. Lindeman, J. E. Visvader, Stem Cells and the Differentiation Hierarchy in Mammary Gland Development. Physiol Rev 100, 489–523 (2020).

15. M. L. Suva, I. Tirosh, Single-Cell RNA Sequencing in Cancer: Lessons Learned and Emerging Challenges. Mol Cell 75, 7–12 (2019).

16. D. Avram et al., Isolation of a novel family of C(2)H(2) zinc finger proteins implicated in transcriptional repression mediated by chicken ovalbumin upstream promoter transcription factor (COUP-TF) orphan nuclear receptors. J Biol Chem 275, 10315–10322 (2000).

17. J. M. Garcia-Aznar, S. Alonso Alvarez, T. Bernal Del Castillo, Pivotal role of BCL11B in the immune, hematopoietic and nervous systems: a review of the BCL11B-associated phenotypes from the genetic perspective. Genes Immun 25, 232–241 (2024).

18. M. J. Lennon, S. P. Jones, M. D. Lovelace, G. J. Guillemin, B. J. Brew, Bcl11b-A Critical Neurodevelopmental Transcription Factor-Roles in Health and Disease. Front Cell Neurosci 11, 89 (2017).

19. S. Cai et al., A Quiescent Bcl11b High Stem Cell Population Is Required for Maintenance of the Mammary Gland. Cell Stem Cell 20, 247–260 e245 (2017).

20. O. Golonzhka et al., Dual role of COUP-TF-interacting protein 2 in epidermal homeostasis and permeability barrier formation. J Invest Dermatol 129, 1459–1470 (2009).

21. V. B. Cismasiu et al., BCL11B functionally associates with the NuRD complex in T lymphocytes to repress targeted promoter. Oncogene 24, 6753–6764 (2005).

22. E. T. Wiles, B. Lui-Sargent, R. Bell, S. L. Lessnick, BCL11B Is Up-Regulated by EWS/FLI and Contributes to the Transformed Phenotype in Ewing Sarcoma. Plos One 8, (2013).

23. G. Ganguli-Indra et al., CTIP2 Expression in Human Head and Neck Squamous Cell Carcinoma Is Linked to Poorly Differentiated Tumor Status. Plos One 4, (2009).

24. H. Abe et al., BCL11B expression in hepatocellular carcinoma relates to chemosensitivity and clinical prognosis. Cancer Med-Us 12, 15650–15663 (2023).

25. G. S. Gulati et al., Single-cell transcriptional diversity is a hallmark of developmental potential. Science 367, 405–411 (2020).

26. S. Z. Wu et al., A single-cell and spatially resolved atlas of human breast cancers. Nat Genet 53, 1334–1347 (2021).

27. T. Kumar et al., A spatially resolved single-cell genomic atlas of the adult human breast. Nature 620, 181–191 (2023).

28. L. Lehtinen et al., PLA2G7 associates with hormone receptor negativity in clinical breast cancer samples and regulates epithelial-mesenchymal transition in cultured breast cancer cells. J Pathol Clin Res 3, 123–138 (2017).

29. Y. Wang, L. Wang, D. Li, H. B. Wang, Q. F. Chen, Mesothelin promotes invasion and metastasis in breast cancer cells. J Int Med Res 40, 2109–2116 (2012).

30. X. Li, Y. Wang, Y. Zhang, B. Liu, Overexpression of MCAM induced by SMYD2-H3K36me2 in breast cancer stem cell properties. Breast Cancer 29, 854–868 (2022).

31. A. Makovec et al., CREB5 as an essential transcription and tumorigenic regulator in basal-like breast and prostate cancer. Journal of Clinical Oncology 42, (2024).

32. C. Hatzis et al., A genomic predictor of response and survival following taxane-anthracycline chemotherapy for invasive breast cancer. JAMA 305, 1873–1881 (2011).

33. S. M. Swain, M. Shastry, E. Hamilton, Targeting HER2-positive breast cancer: advances and future directions. Nat Rev Drug Discov 22, 101–126 (2023).

34. E. Y. Lin et al., Progression to malignancy in the polyoma middle T oncoprotein mouse breast cancer model provides a reliable model for human diseases. Am J Pathol 163, 2113–2126 (2003).

35. K. J. Cheung, E. Gabrielson, Z. Werb, A. J. Ewald, Collective invasion in breast cancer requires a conserved basal epithelial program. Cell 155, 1639–1651 (2013).

36. K. J. Cheung et al., Polyclonal breast cancer metastases arise from collective dissemination of keratin 14-expressing tumor cell clusters. Proc Natl Acad Sci U S A 113, E854–863 (2016).

37. J. Anampa, D. Makower, J. A. Sparano, Progress in adjuvant chemotherapy for breast cancer: an overview. BMC Med 13, 195 (2015).

38. N. Sachs et al., A Living Biobank of Breast Cancer Organoids Captures Disease Heterogeneity. Cell 172, 373–386 e310 (2018).

39. G. Kaur et al., Drug-metabolizing enzymes: role in drug resistance in cancer. Clin Transl Oncol 22, 1667–1680 (2020).

40. A. Bansal, M. C. Simon, Glutathione metabolism in cancer progression and treatment resistance. J Cell Biol 217, 2291–2298 (2018).

41. G. Hu et al., MTDH activation by 8q22 genomic gain promotes chemoresistance and metastasis of poor-prognosis breast cancer. Cancer Cell 15, 9–20 (2009).

42. B. Parajuli, M. L. Fishel, T. D. Hurley, Selective ALDH3A1 inhibition by benzimidazole analogues increase mafosfamide sensitivity in cancer cells. J Med Chem 57, 449–461 (2014).

43. B. Pal et al., Single cell transcriptome atlas of mouse mammary epithelial cells across development. Breast Cancer Res 23, 69 (2021).

44. R. R. Giraddi et al., Single-Cell Transcriptomes Distinguish Stem Cell State Changes and Lineage Specification Programs in Early Mammary Gland Development. Cell Rep 24, 1653–1666 e1657 (2018).

45. R. Liu et al., The prognostic role of a gene signature from tumorigenic breast-cancer cells. N Engl J Med 356, 217–226 (2007).

46. T. Abbas, A. Dutta, p21 in cancer: intricate networks and multiple activities. Nat Rev Cancer 9, 400–414 (2009).

47. B. Mao, Z. Zhang, G. Wang, BTG2: a rising star of tumor suppressors (review). Int J Oncol 46, 459–464 (2015).

48. L. Zhang, J. Yu, B. H. Park, K. W. Kinzler, B. Vogelstein, Role of BAX in the apoptotic response to anticancer agents. Science 290, 989–992 (2000).

49. Y. Ou, S. J. Wang, D. Li, B. Chu, W. Gu, Activation of SAT1 engages polyamine metabolism with p53-mediated ferroptotic responses. Proc Natl Acad Sci U S A 113, E6806–E6812 (2016).

50. M. Diehn et al., Association of reactive oxygen species levels and radioresistance in cancer stem cells. Nature 458, 780–783 (2009).

51. Y. T. Tsai et al., ANGPTL4 Induces TMZ Resistance of Glioblastoma by Promoting Cancer Stemness Enrichment via the EGFR/AKT/4E-BP1 Cascade. Int J Mol Sci 20, (2019).

52. A. Khatib et al., The glutathione peroxidase 8 (GPX8)/IL-6/STAT3 axis is essential in maintaining an aggressive breast cancer phenotype. Proc Natl Acad Sci U S A 117, 21420–21431 (2020).

53. J. Jiang et al., G Protein-Coupled Receptor GPR87 Promotes the Expansion of PDA Stem Cells through Activating JAK2/STAT3. Mol Ther Oncolytics 17, 384–393 (2020).

54. S. Misra et al., Hyaluronan constitutively regulates activation of COX-2-mediated cell survival activity in intestinal epithelial and colon carcinoma cells. J Biol Chem 283, 14335–14344 (2008).

55. S. Long et al., ECM1 regulates the resistance of colorectal cancer to 5-FU treatment by modulating apoptotic cell death and epithelial-mesenchymal transition induction. Front Pharmacol 13, 1005915 (2022).

56. S. L. Huang, C. C. Chao, Silencing of Taxol-Sensitizer Genes in Cancer Cells: Lack of Sensitization Effects. Cancers (Basel) 7, 1052–1071 (2015).

57. C. Lanzi, N. Zaffaroni, G. Cassinelli, Targeting Heparan Sulfate Proteoglycans and their Modifying Enzymes to Enhance Anticancer Chemotherapy Efficacy and Overcome Drug Resistance. Curr Med Chem 24, 2860–2886 (2017).

58. S. Asfaha et al., Krt19(+)/Lgr5(-) Cells Are Radioresistant Cancer-Initiating Stem Cells in the Colon and Intestine. Cell Stem Cell 16, 627–638 (2015).

59. C. Dong, P. Li, Y. Wu, Z. Guo, R. He, The 1q21.3 region driver gene EFNA3 promotes disease progression via inhibition of lung adenocarcinoma cell apoptosis. Transl Cancer Res 11, 1309–1320 (2022).

60. S. T. Susa, A. Hussain, C. V. Preuss, in StatPearls. (Treasure Island (FL), 2025).

61. M. Michael, M. M. Doherty, Tumoral drug metabolism: overview and its implications for cancer therapy. J Clin Oncol 23, 205–229 (2005).

62. A. Singh, J. Settleman, EMT, cancer stem cells and drug resistance: an emerging axis of evil in the war on cancer. Oncogene 29, 4741–4751 (2010).

63. X. Ye et al., Distinct EMT programs control normal mammary stem cells and tumour-initiating cells. Nature 525, 256–260 (2015).

64. C. Kröger et al., Acquisition of a hybrid E/M state is essential for tumorigenicity of basal breast cancer cells (vol 116, pg 7353, 2019). P Natl Acad Sci USA 116, 11553–11554 (2019).

65. L. M. Varela, M. M. Ip, Tumor necrosis factor-alpha: a multifunctional regulator of mammary gland development. Endocrinology 137, 4915–4924 (1996).

66. Y. Wu, B. P. Zhou, TNF-alpha/NF-kappaB/Snail pathway in cancer cell migration and invasion. Br J Cancer 102, 639–644 (2010).

67. M. Endo, T. Masaki, M. Seike, H. Yoshimatsu, TNF-alpha induces hepatic steatosis in mice by enhancing gene expression of sterol regulatory element binding protein-1c (SREBP-1c). Exp Biol Med (Maywood) 232, 614–621 (2007).

68. N. J. Roberts, S. Zhou, L. A. Diaz, Jr., M. Holdhoff, Systemic use of tumor necrosis factor alpha as an anticancer agent. Oncotarget 2, 739–751 (2011).

69. M. Zabala et al., LEFTY1 Is a Dual-SMAD Inhibitor that Promotes Mammary Progenitor Growth and Tumorigenesis. Cell Stem Cell 27, 284–299 e288 (2020).

70. A. Tsherniak et al., Defining a Cancer Dependency Map. Cell 170, 564–576 e516 (2017).

71. W. Yang et al., Genomics of Drug Sensitivity in Cancer (GDSC): a resource for therapeutic biomarker discovery in cancer cells. Nucleic Acids Res 41, D955–961 (2013).

72. J. Puleo, K. Polyak, The MCF10 Model of Breast Tumor Progression. Cancer Res 81, 4183–4185 (2021).

73. S. K. Rehman et al., Colorectal Cancer Cells Enter a Diapause-like DTP State to Survive Chemotherapy. Cell 184, 226–242 e221 (2021).

74. E. Dhimolea et al., An Embryonic Diapause-like Adaptation with Suppressed Myc Activity Enables Tumor Treatment Persistence. Cancer Cell 39, 240–256 e211 (2021).

75. S. Cara, I. F. Tannock, Retreatment of patients with the same chemotherapy: implications for clinical mechanisms of drug resistance. Ann Oncol 12, 23–27 (2001).

76. C. Kadoch et al., Proteomic and bioinformatic analysis of mammalian SWI/SNF complexes identifies extensive roles in human malignancy. Nat Genet 45, 592–601 (2013).

77. G. K. Przybylski, J. Przybylska, Y. Li, Dual role of BCL11B in T-cell malignancies. Blood Sci 6, e00204 (2024).

78. E. Vickridge et al., The function of BCL11B in base excision repair contributes to its dual role as an oncogene and a haplo-insufficient tumor suppressor gene. Nucleic Acids Research 52, 223–242 (2024).

79. M. E. Dourthe et al., The oncogenetic landscape and clinical impact of BCL11B alterations in adult and pediatric T-cell acute lymphoblastic leukemia. Haematologica 108, 3165–3169 (2023).

80. X. B. Zeng et al., Breast cancer stem cells, heterogeneity, targeting therapies and therapeutic implications. Pharmacol Res 163, (2021).

81. C. Verhoef et al., Isolated limb perfusion with melphalan and TNF-alpha in the treatment of extremity sarcoma. Curr Treat Options Oncol 8, 417–427 (2007).

82. H. S. Han et al., Early-Stage Triple-Negative Breast Cancer Journey: Beginning, End, and Everything in Between. Am Soc Clin Oncol Educ Book 43, e390464 (2023).

83. M. Adorno et al., Usp16 contributes to somatic stem-cell defects in Down’s syndrome. Nature 501, 380–384 (2013).

84. D. Kim, J. M. Paggi, C. Park, C. Bennett, S. L. Salzberg, Graph-based genome alignment and genotyping with HISAT2 and HISAT-genotype. Nat Biotechnol 37, 907–915 (2019).

85. S. Anders, P. T. Pyl, W. Huber, HTSeq--a Python framework to work with high-throughput sequencing data. Bioinformatics 31, 166–169 (2015).

86. R. Patro, G. Duggal, M. I. Love, R. A. Irizarry, C. Kingsford, Salmon provides fast and bias-aware quantification of transcript expression. Nat Methods 14, 417–419 (2017).

87. M. I. Love, W. Huber, S. Anders, Moderated estimation of fold change and dispersion for RNA-seq data with DESeq2. Genome Biol 15, 550 (2014).

88. Y. Hao et al., Integrated analysis of multimodal single-cell data. Cell 184, 3573–3587 e3529 (2021).

89. M. Zhao, L. Kong, Y. Liu, H. Qu, dbEMT: an epithelial-mesenchymal transition associated gene resource. Sci Rep 5, 11459 (2015).

90. N. Borcherding et al., Mapping the immune environment in clear cell renal carcinoma by single-cell genomics. Commun Biol 4, 122 (2021).

91. N. Y. Fu et al., EGF-mediated induction of Mcl-1 at the switch to lactation is essential for alveolar cell survival. Nat Cell Biol 17, 365–375 (2015).

92. B. Pal et al., Construction of developmental lineage relationships in the mouse mammary gland by single-cell RNA profiling. Nat Commun 8, 1627 (2017).

93. P. Dalerba et al., Single-cell dissection of transcriptional heterogeneity in human colon tumors. Nat Biotechnol 29, 1120–1127 (2011).

