## Supplementary material for "BCL11B predetermines a persister state in breast cancer that is reversed by TNF*α*": Fig. S1, Fig. S2, Fig. S3, Fig. S4, Fig. S5, Fig. S6, Fig. S7, Fig. S8, Fig. S9, Fig. S10, Fig. S11, Fig. S12

**The PDF file includes:**

Figs. S1 to S12

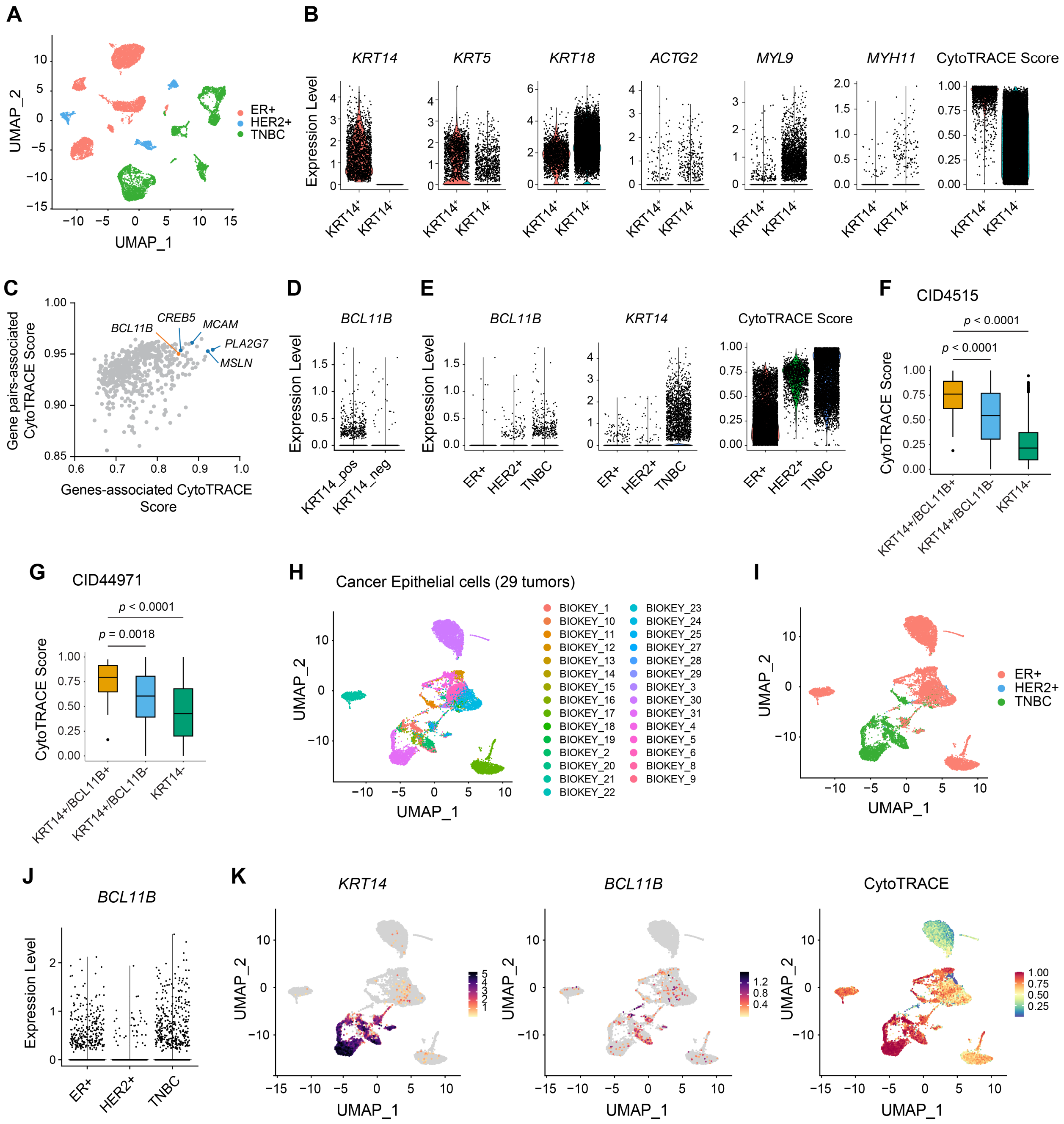


**Fig. S1.** ***BCL11B* represents a distinct marker for less differentiated cancer cells in human breast cancer.** (**A**) UMAP plots showing the cancer epithelial cells colored by tumor subtypes. (**B**) Violin plots comparing the expression levels of the indicated genes between *KRT14*$^{+}$ and *KRT14*$^{-}$ cancer cells. (**C**) Scatterplot showing gene pairs (double positive)-associated CytoTRACE scores (mean) versus genes (single positive)-associated with CytoTRACE scores (mean). (**D**) Violin plots comparing the expression level of *BCL11B* between *KRT14*$^{+}$ and *KRT14*$^{-}$ compartments. Seurat clusters with high *KRT14* expression were designated as the *KRT14*$^{+}$ compartment. (**E**) Violin plots showing the expression levels of the indicated genes across different tumor subtypes. (**F**) Boxplot comparing the levels of CytoTRACE scores among the indicated groups in the cancer epithelial compartment of patient CID4515. (**G**) Boxplot comparing the levels of CytoTRACE scores among the indicated groups in the cancer epithelial compartment of patient CID44971. (**H**) UMAP representation of the cancer epithelial compartment from 29 human breast tumors (public scRNA-seq dataset available at http://biokey.lambrechtslab.org; pre-treatment samples in cohort 1 were analyzed; the metaplastic tumor samples were excluded from the analysis). (**I**) Same UMAP plot colored by tumor subtype. (**J**) Violin plots showing the expression levels of the indicated genes across different tumor subtypes. (**K**) UMAP plots showing the expression patterns of the indicated genes, as well as the levels of CytoTRACE scores.

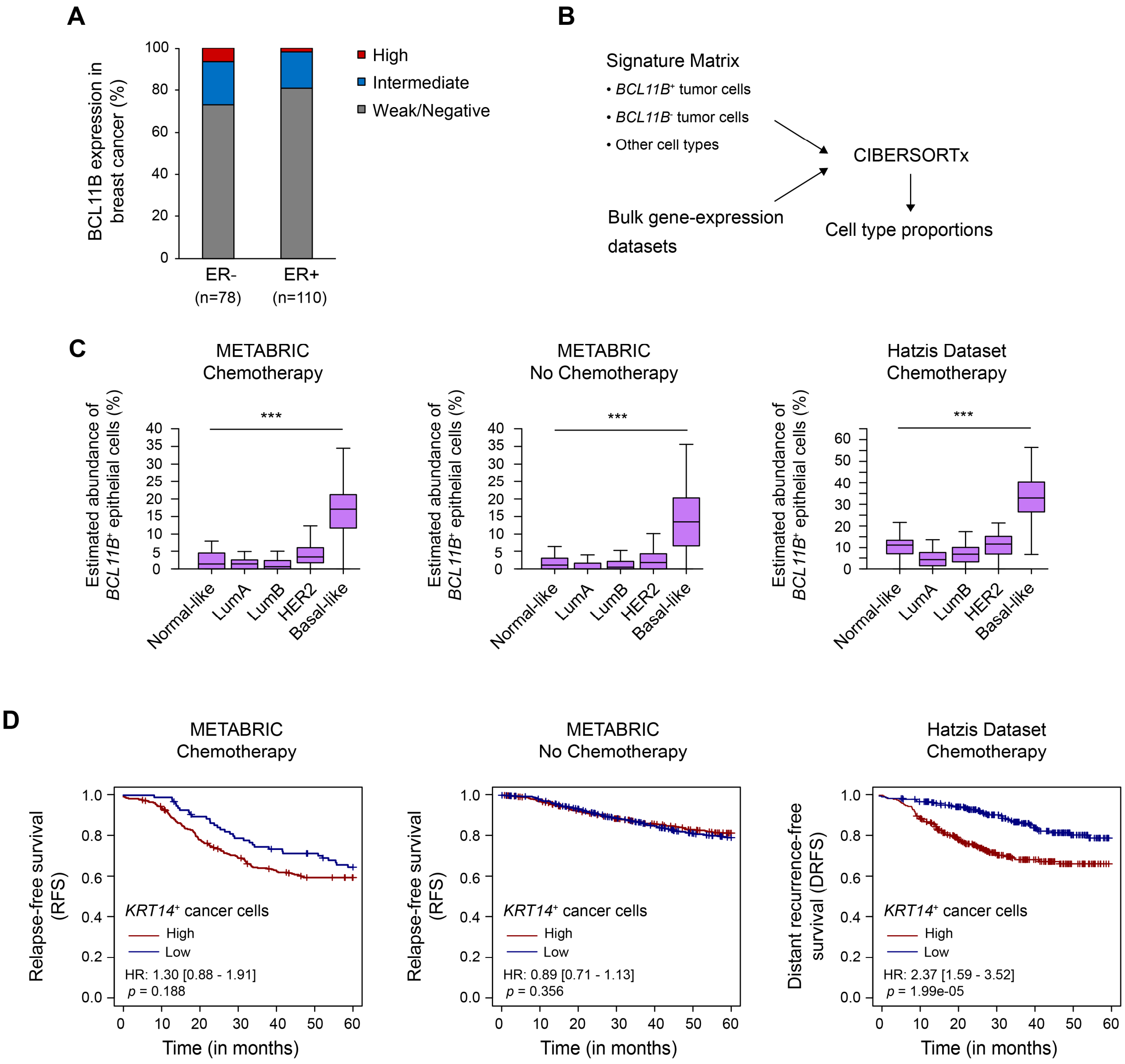


**Fig. S2. *BCL11B*$^+$ cancer cells are associated with chemotherapy resistance in human breast cancer.** (**A**) Quantification of case percentage in each category. High (BCL11B$^+$ cells account for >40% of tumor cells), Intermediate (BCL11B$^+$ cells account for 20%-40% of tumor cells), and Weak/Negative (BCL11B$^+$ cells account for 0-20% of tumor cells). Two breast tissue arrays (BC081116d and BC081116e) were analyzed. (**B**) Schematic diagram showing the strategy of CIBERSORTx-based tumor cell deconvolution. (**C**) Boxplots showing the estimated abundances of *BCL11B*$^+$ tumor cells across the breast cancer subtypes in the METABRIC and Hatzis datasets. (**D**) Kaplan-Meier curve showing differences in relapse-free survival stratified by the median abundance of *KRT14*$^+$ cancer epithelial cells in the METABRIC cohort (n = 392, chemotherapy; n = 1,503, no chemotherapy) and in the Hatzis cohort (n = 508).

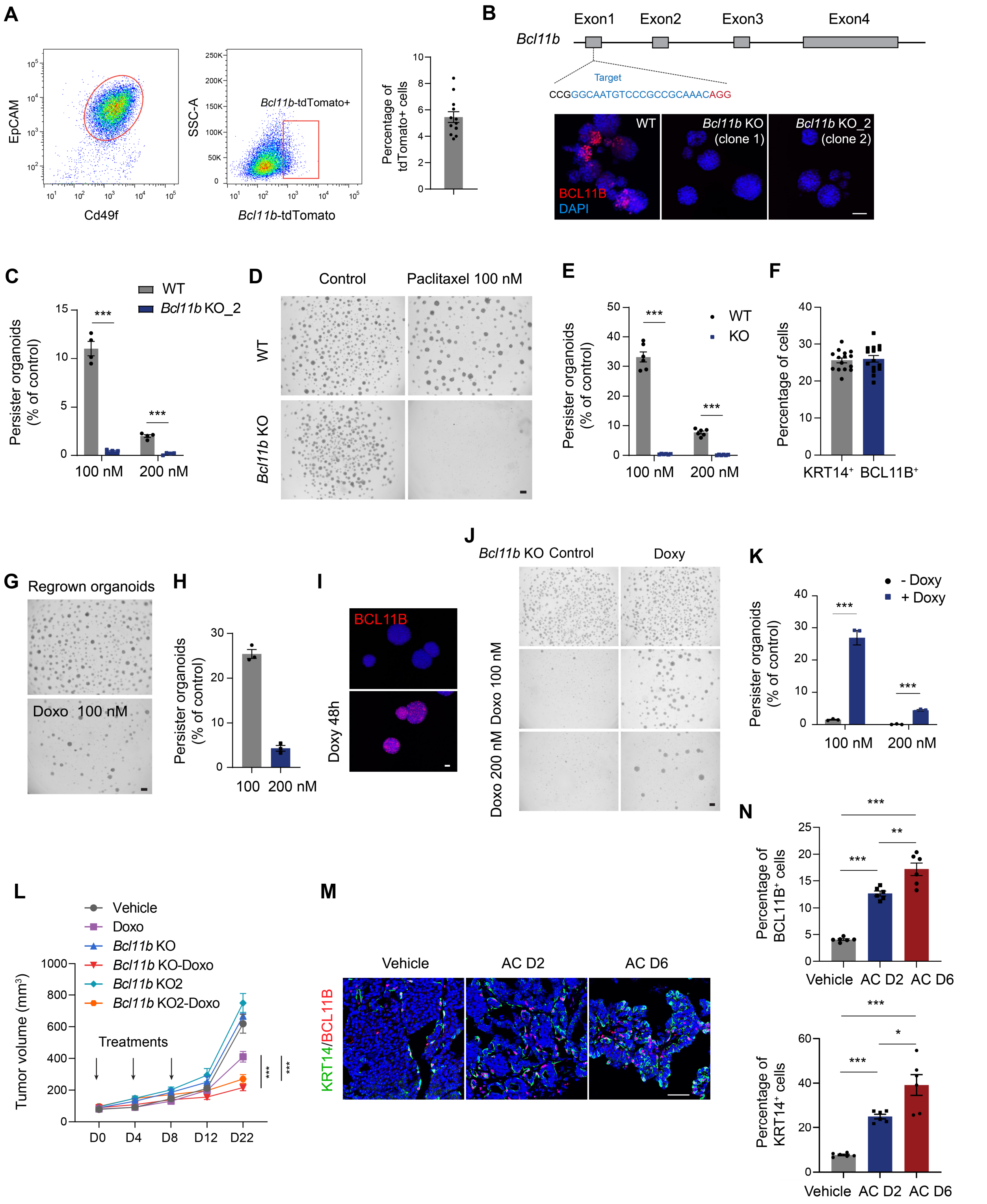


**Fig. S3. BCL11B$^{+}$ tumor cells demonstrate multi-drug resistance ability.** (**A**) Representative FACS plots of tumor cells from PyMT-*Bcl11b*$^{\mathrm{tdTomato}}$ mice. Quantification of the percentage of tdTomato+ cells is shown; n = 12 tumor samples. (**B**) Upper panel: A schematic diagram of

CRISPR/Cas9-mediated *Bcl11b* knockout targeting strategy. Lower panel: BCL11B immunostaining confirmed the knockout of *Bcl11b* in two single clones. (**C**) Quantification of the frequency of drug-resistant persister organoids in the indicated conditions; n = 4 cell culture wells from 3 experiments. (**D**) Tumor organoid formation in WT or *Bcl11b* knockout group after paclitaxel treatment (100 or 200 nM). Scale bar, 300 µm. (**E**) Quantification of the frequency of drug-resistant persister organoids under indicated conditions; n = 6 cell culture wells from 3 experiments. (**F**) Percentage of $KRT14^+$ and $BCL11B^+$ cells in the regrown organoids; n = 15 organoids from 3 experiments. (**G**) Drug treatment assay in the regrown tumor organoids. Scale bar, 300 µm. (**H**) Quantification of the frequency of drug-resistant persister organoids from regrown organoid cells under indicated conditions; n = 3 experiments. (**I**) An inducible *Bcl11b* over-expression vector was introduced to the *Bcl11b* knockout organoids via lentivirus. Representative images of BCL11B immunostaining in control organoids and doxycycline (Doxy, 100 ng/mL) treated organoids at 48 hours are shown. Scale bar, 50 µm. (**J**) *Bcl11b*-KO cells with an inducible *Bcl11b* overexpressing construct were treated with or without doxycycline for three days and then dissociated into single cells. The single cells were exposed to either doxorubicin or vehicle for 24 hours, followed by drug washout and recovery. Doxycycline was provided to the indicated groups throughout the experiment. Scale bar, 300 µm. (**K**) Frequency of persister organoids derived from WT or *Bcl11b* overexpression cells under drug treatment; n = 3 experiments. (**L**) Tumor growth curves showing WT, *Bcl11b* KO, *Bcl11b* KO_2 tumor growth upon saline (Ctrl) treatment, or doxorubicin treatment (1 mg/kg). n = 5 mice per group, two-way ANOVA. (**M**) Representative images of KRT14 and BCL11B immunostaining in untreated or AC (doxorubicin + cyclophosphamide)-treated PyMT tumor samples (2 days or 6 days after the final treatment). (**N**) Percentage of $KRT14^+$ and $BCL11B^+$ cells in WT tumor under indicated conditions; n = 6 tumors. Nuclei were counter-stained with DAPI (blue). Data are presented as mean ± SEM. Scale bar, 50 µm. *$p < 0.05$, **$p < 0.01$, ***$p < 0.001$.

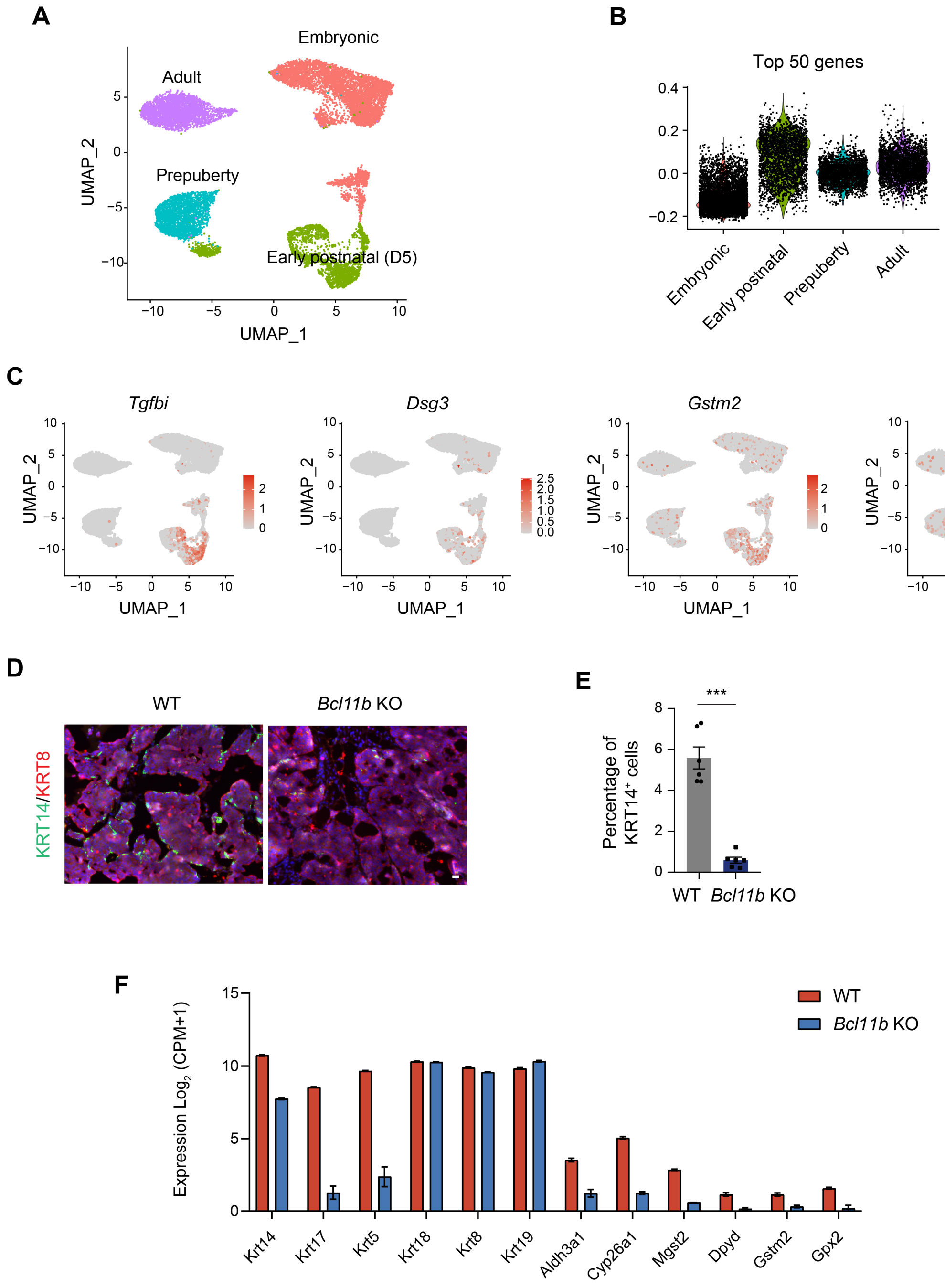


**Fig. S4. The *Bcl11b*$^{+}$ cancer cell population is linked to a rudimentary basal-like cell state during development with an intrinsic drug resistance program.** (**A**) UMAP plot of *Krt14*-positive basal and basal-like cells across 4 different developmental stages of the mammary gland (GEO: GSE164017). The *Krt14*-positive cell clusters from the different stages were combined and analyzed together. (**B**) Violin plots showing the expression score of the top 50 *Bcl11b* co-expressed genes in tumors within the basal and basal-like mammary cells across developmental stages. (**C**) UMAP plots displaying the expression patterns of indicated genes in the scRNA-seq dataset of mammary basal and basal-like cells. (**D**) Immunofluorescence staining of orthotopic tumors derived from WT or *Bcl11b* knockout organoid cells. Nuclei were counter-stained with DAPI (blue). Scale bar, 50 μm. (**E**) Percentage of KRT14$^+$ cells in WT and *Bcl11b* KO tumors; n = 6 tumors. (**F**) Expression levels of the basal cell markers, luminal cell markers, and selected drug resistance genes in the bulk RNA-seq dataset of *Bcl11b* WT and knockout cells. Data are presented as mean ± SEM. *$p < 0.05$, **$p < 0.01$, ***$p < 0.001$.

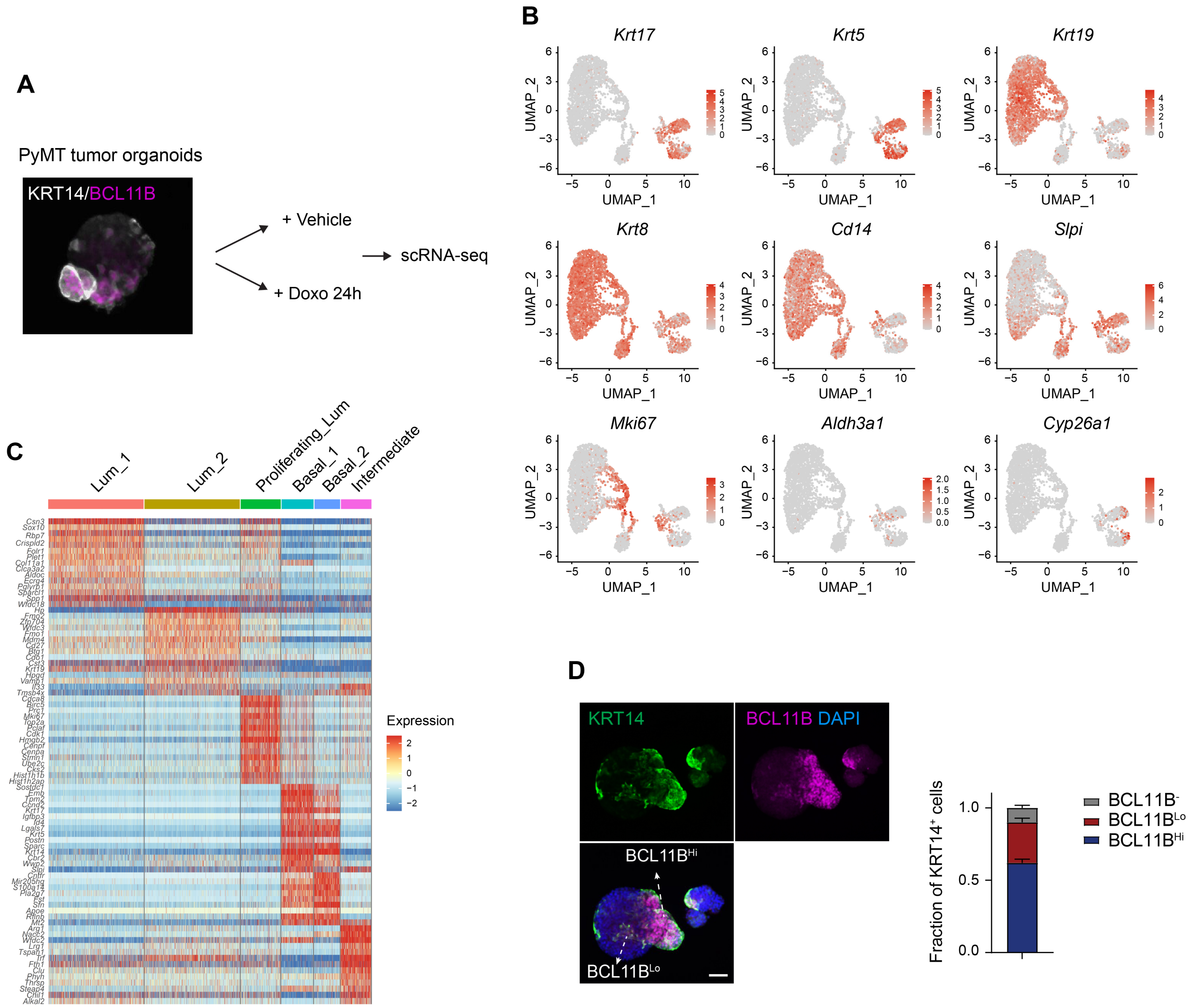


**Fig. S5. Single-cell RNA-seq analysis of treatment-naïve tumor organoids.** (**A**) Schematic diagram showing the scRNA-seq experimental design. (**B**) UMAP plots showing the expression levels of selected marker genes in the vehicle-treated control tumor cells. (**C**) Heatmap showing top marker genes for each cluster in the control dataset. (**D**) Quantifications of the percentage of $BCL11B^{Hi}$, $BCL11B^{Lo}$ and $BCL11B^{-}$ cells within the $KRT14^{+}$ cell population in tumor organoids; n = 7 tumor organoids. Data are presented as mean ± SEM. The representative immunostaining image of tumor organoids is shown on the left. KRT14, green; BCL11B, purple. Nuclei were counter-stained with DAPI (blue). Scale bar, 50 μm.

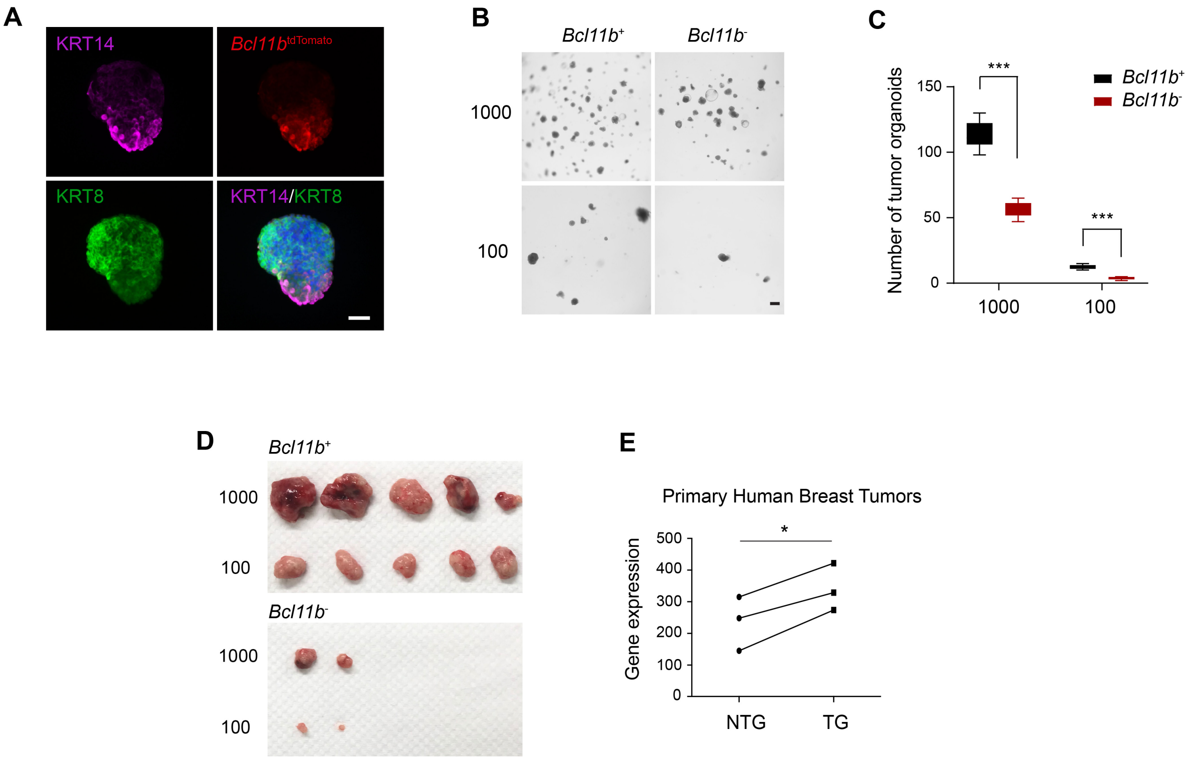


**Fig. S6. *Bcl11b*$^{+}$ tumor cells maintain the differentiation potential and display a higher tumorigenic ability.** (**A**) Representative images of *Bcl11b*$^{tdTomato}$ immunofluorescence, KRT14, and KRT8 staining in tumor organoids grown from FACS-sorted single *Bcl11b*$^{tdTomato}$ positive cells. Scale bar, 50 µm. (**B**) Representative images of tumor organoids generated from the indicated number of *Bcl11b*$^{tdTomato}$ positive or *Bcl11b*$^{tdTomato}$ negative tumor cells. (**C**) Quantification of the number of organoids generated from the indicated number of cells; n = 3 experiments. (**D**) Photographs of representative tumors from mice engrafted with the indicated number of *Bcl11b*$^{tdTomato}$ positive or *Bcl11b*$^{tdTomato}$ negative tumor cells. (**E**) Comparison of *BCL11B* expression in non-tumorigenic and tumorigenic samples of human breast tumors (GEO: GSE6883). Nuclei were counter-stained with DAPI (blue). Data are presented as mean ± SEM; n = 3. $*p < 0.05$, $**p < 0.01$, $***p < 0.001$.

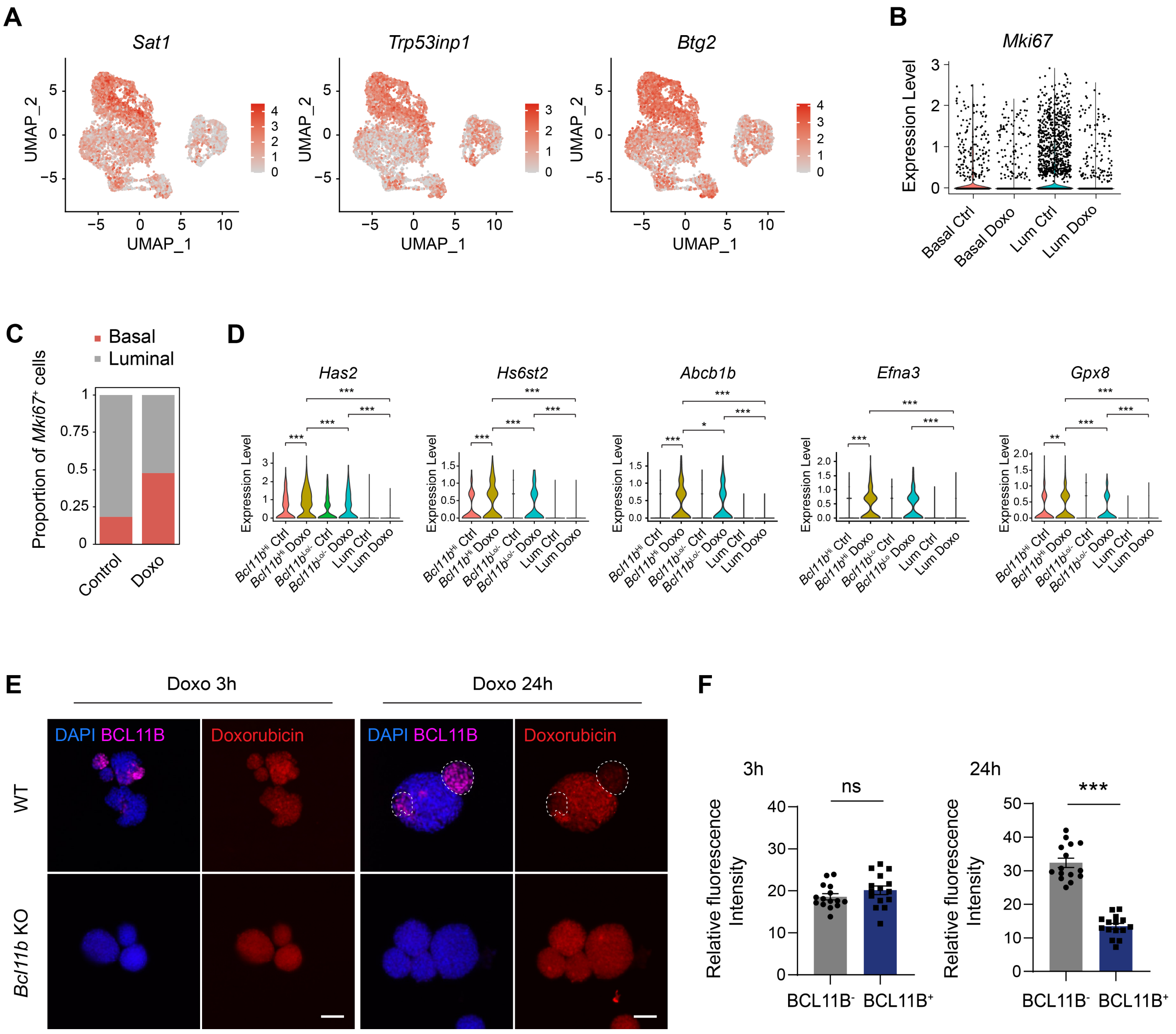


**Fig. S7. *Bcl11b*$^{+}$ basal-like cells are tolerant to chemotherapy treatment.** (**A**) UMAP plots showing the expression levels of cell cycle arrest and apoptosis-associated genes in the combined dataset. (**B**) Violin plots showing the changes in *Mki67* expression in the basal and luminal compartments upon the drug treatment. (**C**) Relative proportions of *Mki67*+ basal and luminal tumor cells in the untreated and drug-treated tumor organoids. (**D**) Violin plots displaying the expression of representative drug-resistance related genes across the six subsets. (**E**) WT or *Bcl11b*-deficient tumor organoids were treated with doxorubicin (100 nM) for 3 hours and 24 hours. Representative images of BCL11B immunostaining and doxorubicin fluorescence at different time points are shown. Doxorubicin autofluorescence can be used to measure the intracellular drug concentration. BCL11B$^{+}$ cell clusters at 24 h were highlighted with circles. Scale bar, 50 μm. (**F**) The fluorescence intensity of doxorubicin was quantified in BCL11B$^{-}$ and BCL11B$^{+}$ cells after 3, 24 or 48-hour treatment; n = 15 organoids from 3 experiments. Nuclei were counter-stained with DAPI (blue). Data are presented as mean ± SEM. *p < 0.05, **p < 0.01, ***p < 0.001.

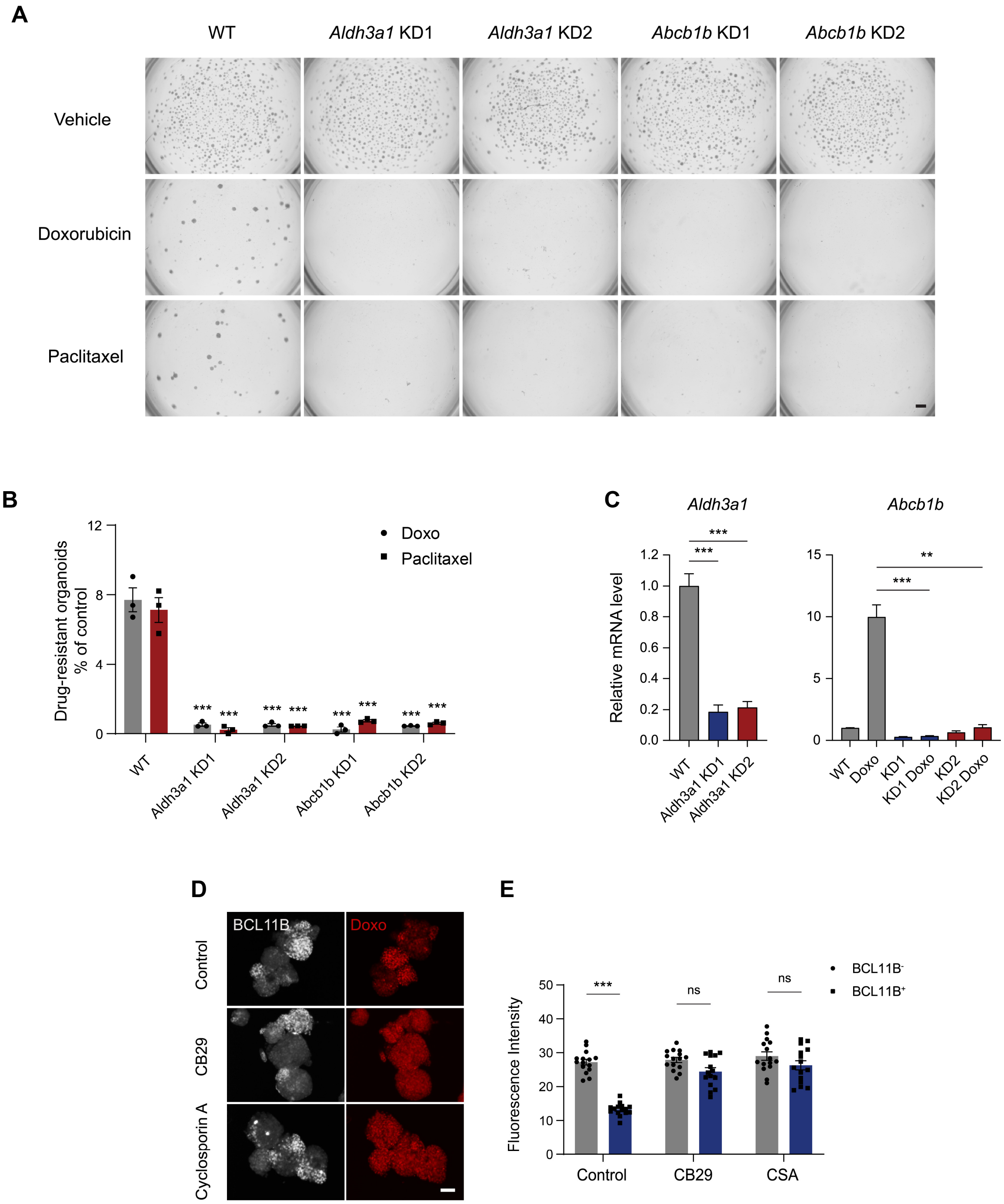


**Fig. S8. The drug detoxification program contributes to drug resistance in BCL11B$^+$ cells.** (**A**) *Ex vivo* drug treatment assay was performed to assess the drug sensitivity of WT, *Aldh3a1*, and *Abcb1b* knockdown tumor cells. Scale bar, 500 µm. (**B**) Quantification of the percentage of drug-resistant organoids under the indicated conditions; n = 3 experiments. Data are presented as mean ± SEM. (**C**) Real-time PCR analysis of indicated genes in the indicated groups with *Actb* as an internal control; n = 3 experiments. (**D**) Tumor organoids were either left untreated or pre-treated with the Aldh3a1 inhibitor (CB29, 70 µM) or Abcb1b inhibitor (Cyclosporin A (CSA), 5 μM) for 24 hours before exposure to a 24-hour drug treatment (doxorubicin, 100 nM). Representative images of BCL11B immunostaining and doxorubicin fluorescence are shown. Scale bar, 50 μm. (**E**) Quantification of the fluorescence intensity of doxorubicin in $BCL11B^-$ and $BCL11B^+$ cells after 24-hour drug treatment in the absence or presence of indicated inhibitors; n = 15 organoids from 3 experiments. Data are presented as mean ± SEM. *p < 0.05, **p < 0.01, ***p < 0.001.

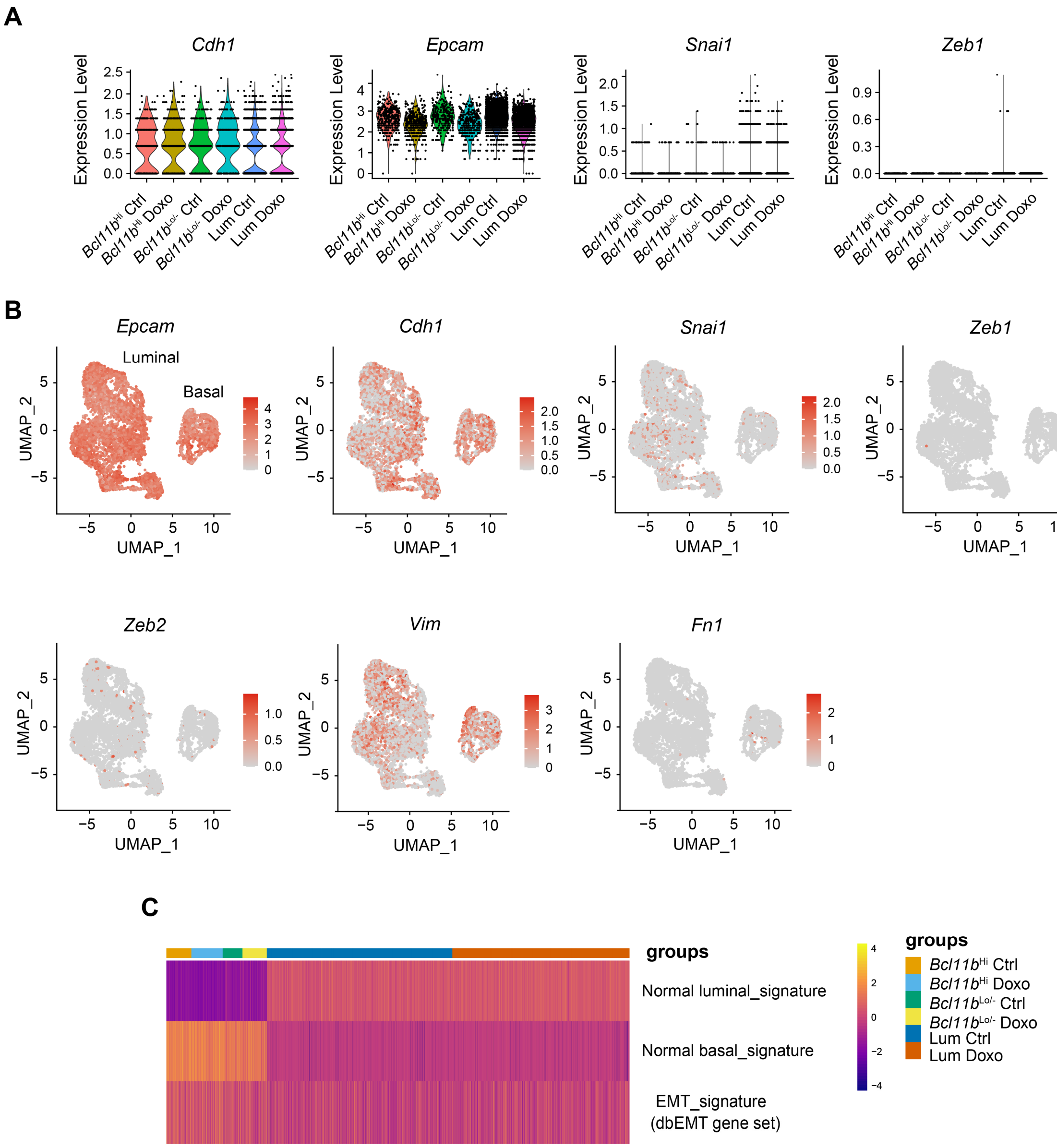


**Fig. S9. *Bcl11b*$^{+}$ basal-like cells are distinct from cells with EMT activation.** (**A**) Violin plots displaying the expression of indicated marker genes across the six subsets. (**B**) UMAP plots showing the expression levels of EMT-related genes in the combined dataset. (**C**) Enrichment analysis of the Normal luminal signature, Normal basal signature, and EMT signature (dbEMT gene set) in indicated subsets of the tumor cells.

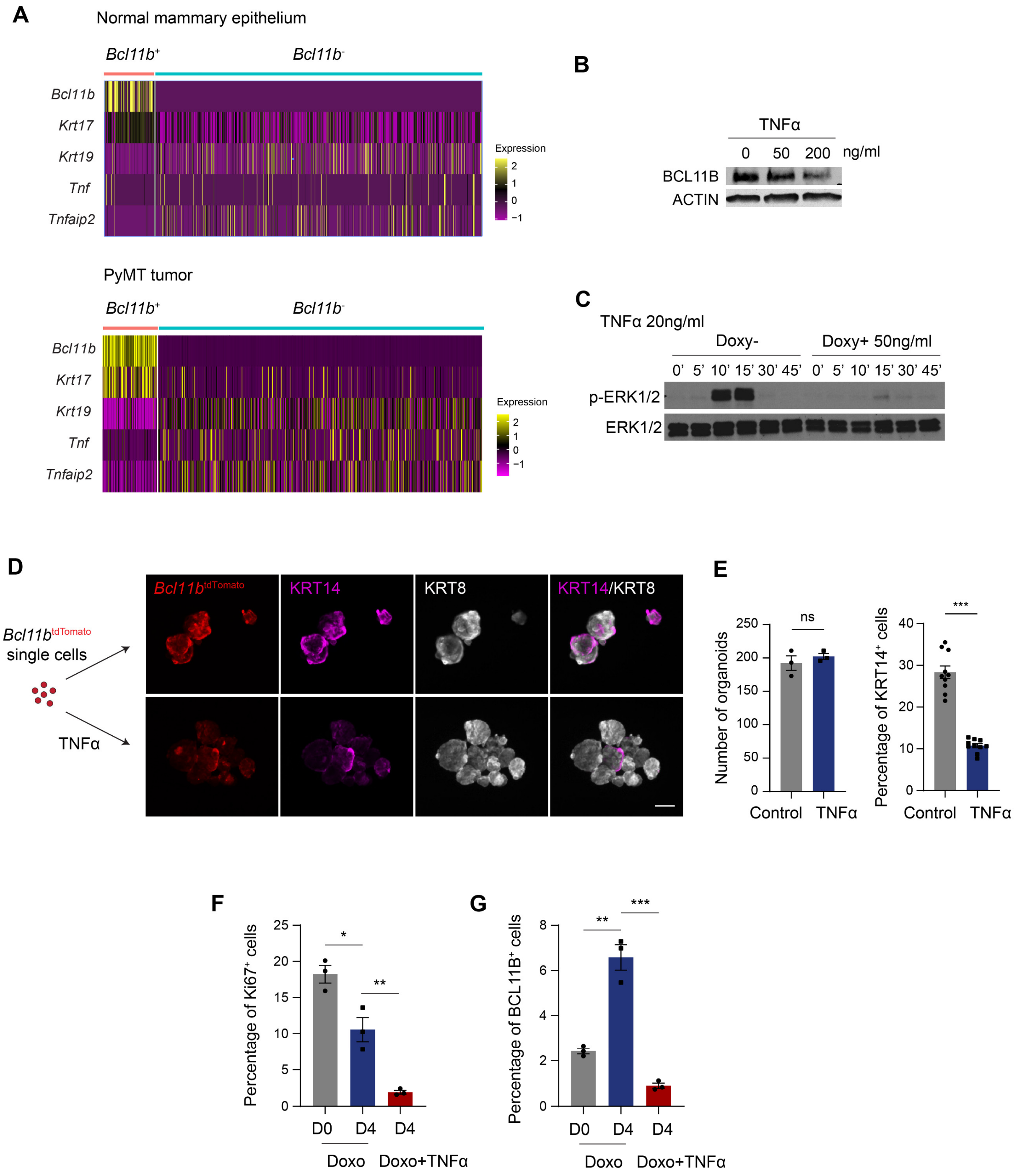


**Fig. S10. TNFα treatment prevents the emergence of cancer persisters by disrupting BCL11B expression.** (**A**) Upper panel: heatmap displaying the expression pattern of selected genes in *Bcl11b*+ and *Bcl11b*- cells in the scRNA-seq dataset of normal mouse mammary epithelial cells (GEO: GSE109774). Lower panel: heatmap displaying the expression pattern of selected genes in *Bcl11b*+ and *Bcl11b*- cells in the single-cell RNA-seq data of PyMT organoids. (**B**) Western blot analysis of BCL11B expression upon treatment of TNFα. (**C**) Overexpression of BCL11B inhibits TNFα-induced ERK phosphorylation. WB analysis of the level of p-ERK1/2 upon TNFα treatment in the absence or presence of doxycycline is shown. (**D**) Immunofluorescence analysis of KRT14, KRT8, and *Bcl11b*-tdTomato in the tumor organoids derived from single *Bcl11b*$^{tdTomato}$ positive cells in the absence or presence of TNFα treatment. Scale bar, 50 μm. (**E**) Quantification of the percentage of KRT14$^{+}$ cells in the tumor organoids derived from single *Bcl11b*$^{tdTomato}$ positive cells under different conditions; n = 10 organoids from 3 experiments. (**F**) Quantification of the percentage of Ki67$^{+}$ cells in indicated groups; n=3 tumors. (**G**) Quantification of the percentage of BCL11B$^{+}$ cells in indicated groups; n=3 tumors. Data are presented as mean ± SEM. *$p < 0.05$, **$p < 0.01$, ***$p < 0.001$.

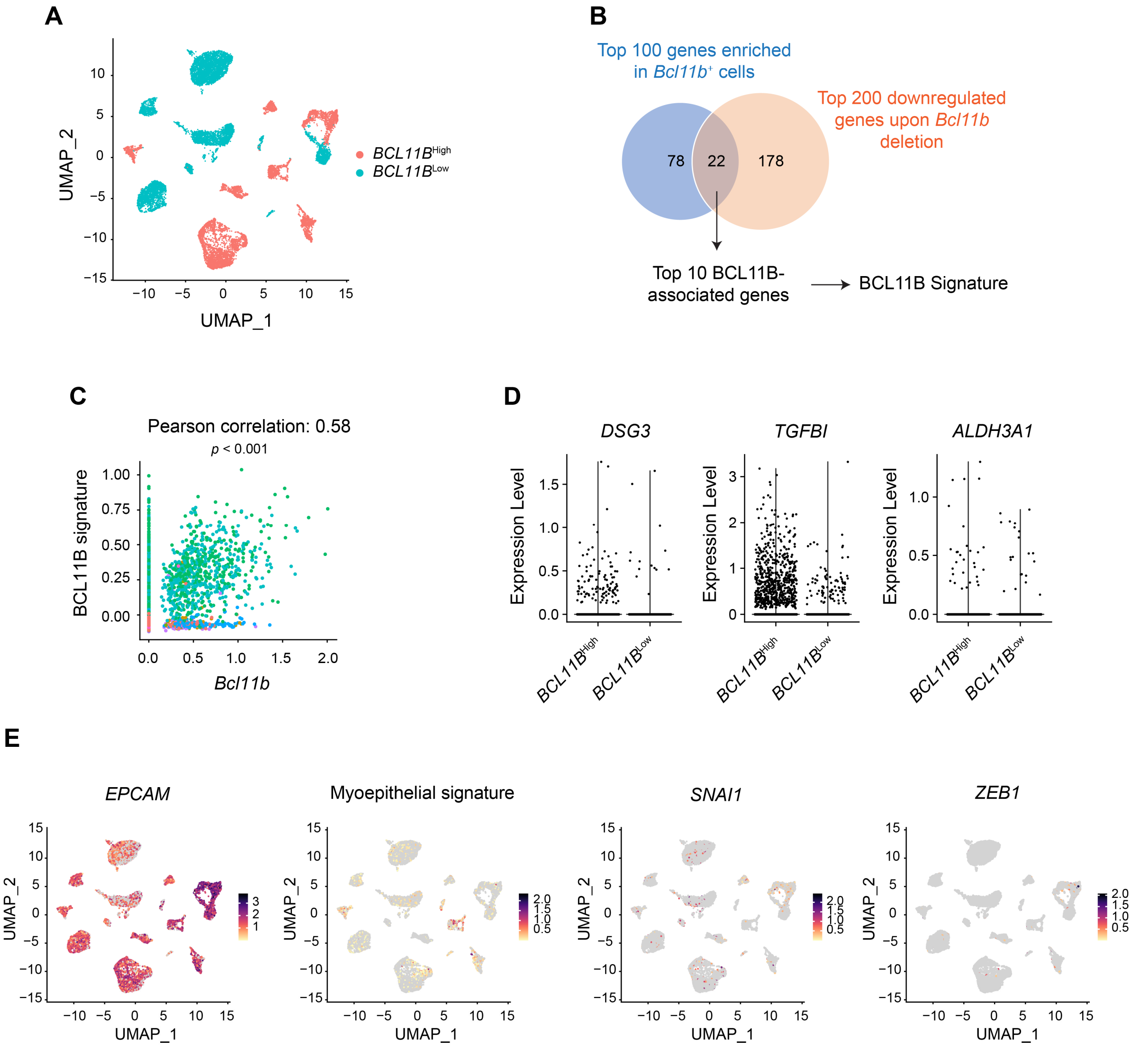


**Fig. S11. The BCL11B$^{+}$ cell state is conserved in human breast cancer.** (**A**) UMAP plots showing the cancer epithelial cells in the single-cell atlas colored by *BCL11B*-high and *BCL11B*-low groups. (**B**) Schematic diagram showing the strategy of generating the 'BCL11B signature' based on bulk RNA-seq analysis. (**C**) Correlation between *Bcl11b* and the BCL11B signature in the scRNA-seq data of tumor organoids. (**D**) Violet plots showing the expression of selected genes in the *BCL11B*-high and *BCL11B*-low compartments. (**E**) UMAP plots displaying the expression patterns of *EPCAM*, the myoepithelial signature (*ACTA2*, *ACTG2*, *MYH11*, *MYL9*, *TAGLN*, and *MYLK*), *SNAI1*, and *ZEB1* in the epithelial cell compartment of human breast cancer.

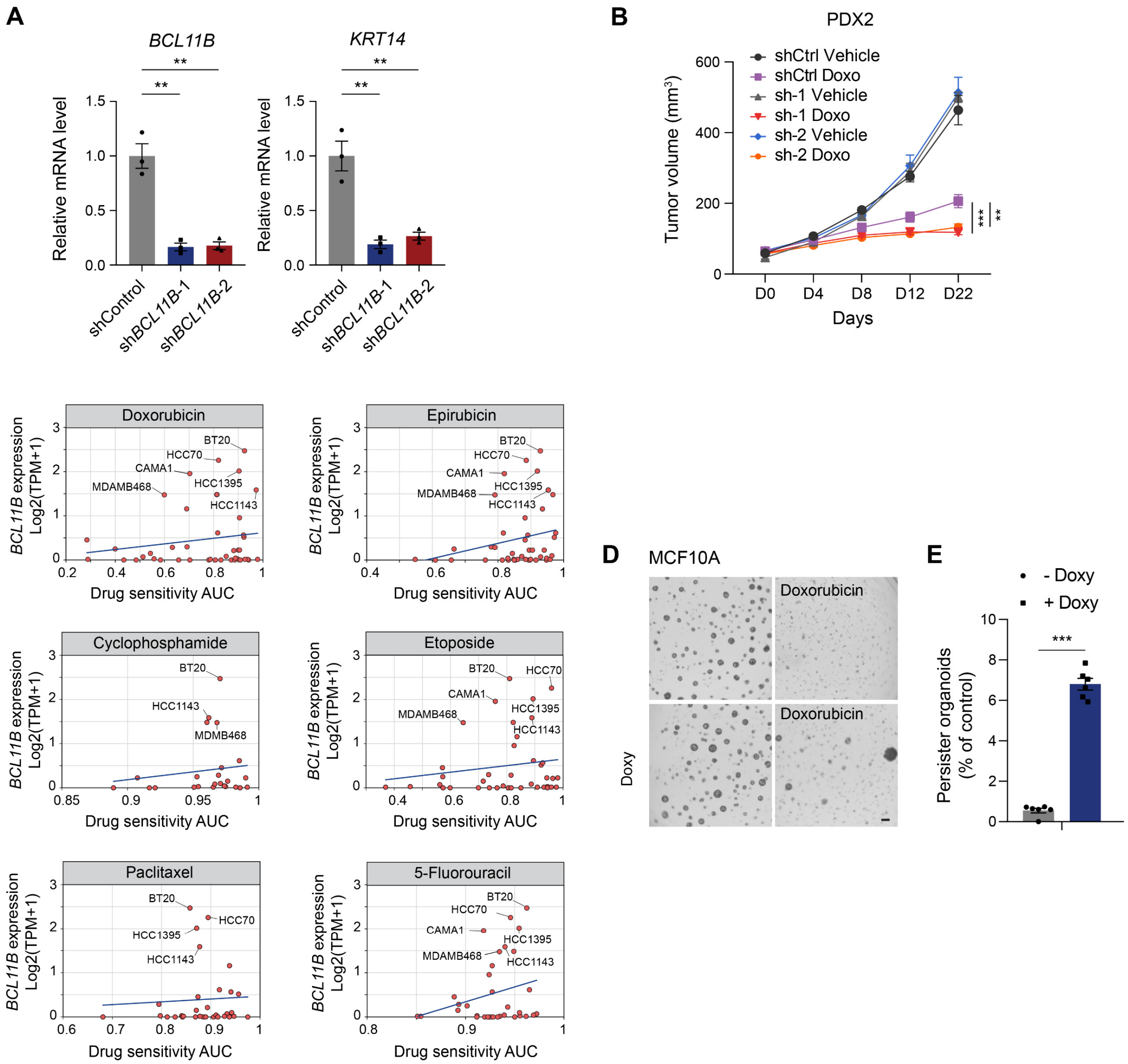


**Fig. S12. BCL11B promotes treatment resistance in human breast cancer.** (**A**) Real-time PCR quantification of *BCL11B* and *KRT14* mRNA levels in control PDX2 and *BCL11B* knockdown PDX2 cells. n = 3 experiments. (**B**) Tumor growth curves showing the growth of WT PDX2 tumors or *BCL11B* knockdown PDX2 tumors upon saline (Vehicle) and doxorubicin (Doxo) treatment. n = 6 mice per group, two-way ANOVA. (**C**) *BCL11B* expression in breast cancer cell lines versus drug sensitivity AUC for chemotherapeutic drugs based on the data from the DepMap and GDSC databases. A higher AUC value implies less drug sensitivity. (**D**) Drug treatment assay in control MCF10A cells and BCL11B over-expressing MCF10A cells. MCF10A cells with an inducible BCL11B overexpressing construct were treated with or without doxycycline for three days and then dissociated into single cells. The single cells were cultured in Matrigel in the presence or absence of doxorubicin for 24 hours, followed by drug washout and recovery. Doxycycline was provided to the indicated groups for the duration of the experiment. Scale bar, 300 μm. (**E**) Quantification of the frequency of drug-resistant organoids in the indicated conditions; n = 6 cell culture wells from 3 experiments. Data are presented as mean ± SEM. *$p < 0.05$, **$p < 0.01$, ***$p < 0.001$.
